# Ciliary tip actin dynamics regulate the cadence of photoreceptor disc formation

**DOI:** 10.1101/2022.11.10.516020

**Authors:** Roly Megaw, Abigail Moye, Zhixian Zhang, Fay Newton, Fraser McPhie, Laura C. Murphy, Lisa McKie, Feng He, Melissa K. Jungnickel, Alex von Kriegsheim, Laura M. Machesky, Theodore Wensel, Pleasantine Mill

## Abstract

As signalling organelles, primary cilia regulate their membrane G protein-coupled receptor (GPCR) content by ectocytosis, a process requiring localised actin dynamics at their tip to alter membrane shape.(1, 2) Mammalian photoreceptor outer segments comprise an expanse of folded membranes (discs) at the tip of highly-specialised connecting cilia (CC), in which photosensitive GPCRs like rhodopsin are concentrated. In an extraordinary feat of biology, outer segment discs are shed and remade daily.(3) Defects in this process, due to genetic mutations, cause retinitis pigmentosa (RP), an untreatable, blinding disease. The mechanism by which photoreceptor cilia generate outer segments is therefore fundamental for vision yet poorly understood. Here, we show the membrane deformation required for outer segment disc genesis is driven by dynamic changes in the actin cytoskeleton in a process akin to ectocytosis. Further, we show *RPGR*, a leading causal RP gene, regulates activity of actin binding proteins crucial to this process. Disc genesis is compromised in *Rpgr* mouse models, slowing the actin dynamics required for timely disc formation, leading to aborted membrane shedding as ectosome-like vesicles, photoreceptor death and visual loss. Manipulation of actin dynamics partially rescues the phenotype, suggesting this pathway could be targeted therapeutically. These findings help define how actin-mediated dynamics control outer segment turnover.

## Main

Most mammalian cells assemble a primary cilium; a microtubule-based structure that protrudes from the cell body and functions as a sensory organelle by detecting changes in the extracellular environment and initiating signalling.(4) Cilia dysfunction, due to pathogenic mutations in critical genes, leads to a spectrum of human diseases termed the ciliopathies, which comprise multisystem disorders of the brain, lung, kidney and eye, amongst others.(4) Thus, tight control of cilia signalling is crucial for human health.

Cilia function is optimised by compartmentalising the initiators of signalling cascades, such as G protein-coupled receptors (GPCRs), in its membrane. This is achieved by high volume trafficking to the cilia, but more recently it has been shown that dynamic membrane changes at the ciliary tip can regulate GPCR concentration within the cilium in a process termed ectocytosis, which involves the shedding of cilia-membrane-derived vesicles into the extracellular space.(1, 2) Ectosome formation is facilitated by local changes in the actin cytoskeleton to initiate the membrane deformation required to form these structures that will be subsequently pinched off and shed. How important this biological process is across cell types and in the context of human health remains unclear.

The photoreceptor contains one of the most highly specialised primary cilia, that has evolved to optimise our visual processing capabilities by compartmentalising its photosensitive GPCRs within hundreds of disc-like membranous processes that stack on top of each other at the distal end of the cilia to form the cell’s outer segment (OS).(5, 6) To enable recycling of its contents, the photoreceptor OS is completely renewed every 7 to 10 days,(3) with distal discs shed for phagocytosis by the underlying retinal pigment epithelium (RPE).(7, 8) A fine balance, with continuous birth of photoreceptor discs to replace the shed OS material, is critical to support vision. The mechanism that drives the ciliary membrane remodelling required for disc formation is yet to be fully determined, but evidence is mounting that it is an actin-driven process(8–11) and it has been speculated that the process has evolved as a form of ectocytosis.(10, 12, 13) Failure to renew photoreceptor discs has been implicated in retinitis pigmentosa (RP),(13) a heterogenous group of inherited retinal dystrophies affecting 1 in 3000 people(14) that cause blindness. Patients present with night blindness and progressive constriction of their visual fields, prior to loss of central vision, as their photoreceptors degenerate.

Here, using cryo-electron tomography, novel reporter mice and advanced live imaging techniques, we provide evidence supporting a model whereby the membrane deformation required for photoreceptor disc formation is an actin driven process akin to ectocytosis. Further, we show that retinitis pigmentosa GTPase regulator *(RPGR)*, mutations in which cause 15% of RP,(15) functions to control the tempo of disc formation by binding the actin severing protein cofilin in the distal photoreceptor cilia and regulating its activity. *RPGR* mutations compromise cofilin activity, resulting in lengthened actin bundles in the newly forming disc, slowing their maturation. As a result, compromised discs are instead shed as ectosome-like vesicles, resulting in a reduced rate of disc morphogenesis, outer segment abnormalities, retinal stress, photoreceptor degeneration and loss of vision. We conclude, therefore, that highly regulated actin control in the nascent photoreceptor disc controls the cadence of disc genesis in the same manner as ectosome formation and that *Rpgr* plays a crucial role in the process.

### Disc formation is an active, actin driven process

There is growing evidence that disc formation is actin-dependent.(8–11) It has not been definitively shown, however, if this is due to an active process, whereby progressive nucleation of actin microfilaments (akin to lamellipodia formation)(16) mechanically deform the membrane. Alternatively, it could be a passive process (such as blebbing), whereby loss of adhesion between the membrane and underlying cortical actin allows the hydrostatic pressure within a cell to deform the membrane.(17) To distinguish between these possibilities in situ at nanoscale, we used cryo-electron tomography (cryoET).(18) Flash freezing isolated mouse rod photoreceptor outer segments (ROS) allows visualisation of 3-dimensional architecture at an ultrastructural level by creating 3-dimensional maps from a tilt series of electron tomograph images. Three dimensional reconstructions showed microfilaments extending from the distal CC into the nascent disc (**Fig. 1a-c**). Sub-tomogram averaging of these microfilaments revealed similarities to filamentous actin’s published structure of 166.67° twists per molecule and a 27.8 Å rise per molecule (**Fig. 1d-f**).(19) Using live imaging of retinal slice cultures incubated with silicon rhodamine-actin (SiR-Actin), a fluorogenic, cell permeable probe based on the highly specific F actin binding drug jasplakinolide, we observed elongation, constriction and movement of the microfilaments within the connecting cilium evident over time (**Extended Data Movie 1**). We thus conclude that dynamic actin changes occur within nascent photoreceptor discs, suggesting a role for actin in disc genesis.

**Fig. 1.**
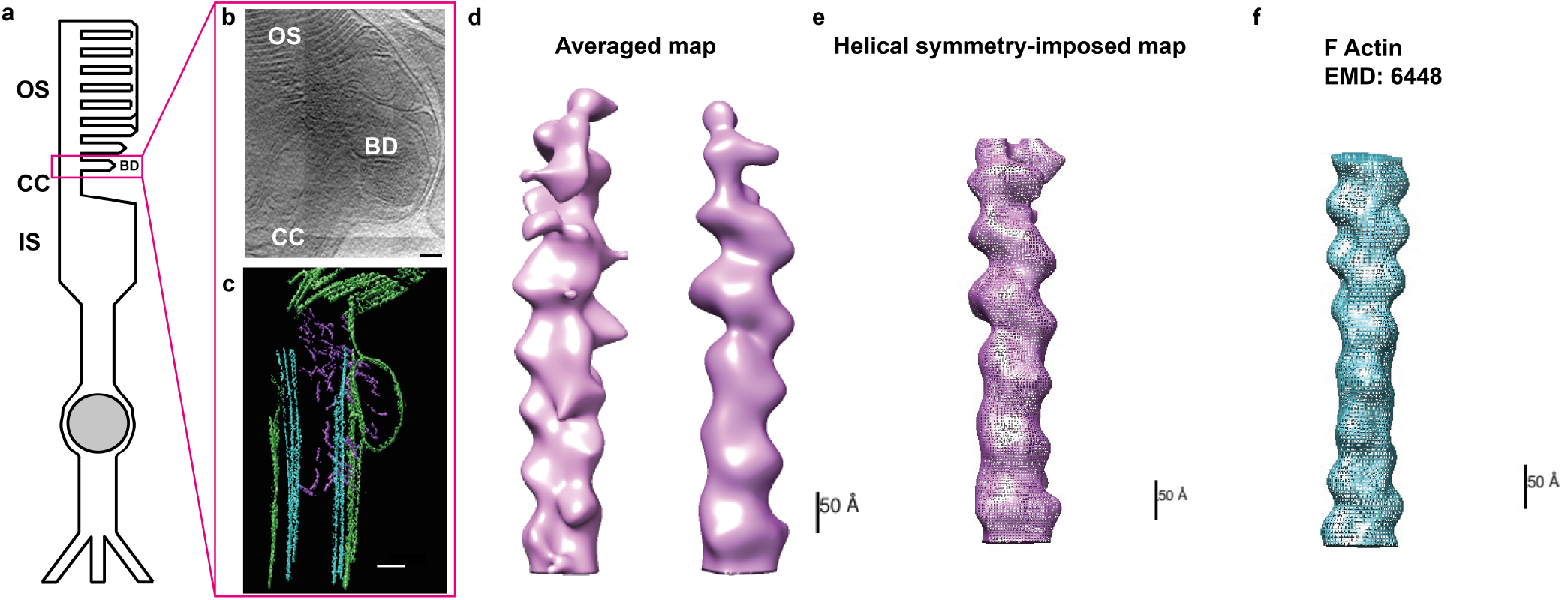
Photoreceptor disc formation is an actin-driven process. **(a)** Cartoon schematic of a photoreceptor depicts the cell’s inner segment (IS) with the highly modified connecting cilium (CC) extending from its apex. From here an expanse of folded membrane extends, forming the outer segment (OS) discs. Basal discs (BD) are continually added at its base. **(b)** A projection from a tomogram of the basal disc of a wild type photoreceptor. **(c)** A slice through a tomographic reconstruction, segmented to highlight the ciliary and disc membranes (green), microtubule based axoneme (cyan) and microfilaments (purple). Microfilaments extend from the connecting cilium into the basal disc. **(d)** Front and back side impression of a subtomogram averaged map of microfilaments extending into the BD, with helical symmetry-imposed map **(e). (f)** Previously published structure of F-actin (EMBD-6448), highlighting 166.67 degree twist per molecule and 27.8 Å rise per molecule. (Scale bars; b,c = 100 nm; d-f = 50 Å)

### RPGR is required for photoreceptor outer segment turnover

Mutations in *RPGR* account for 70-90% of X-linked RP (XLRP) and 10-15% of all RP and result in a severe form of disease.(15) RPGR is alternatively spliced, with a retinal specific isoform containing a repetitive, disordered C-terminal domain of unknown function (conventionally termed ‘ORF15’) that is a mutational hotspot for disease.(20) RPGR localises to the photoreceptor CC and its loss results in perturbation of CC actin regulation, with subsequent photoreceptor degeneration.(12, 21–23) To determine if RPGR plays a role in actin-driven disc genesis, we generated novel *Rpgr*-mutant mice harbouring humanised disease-causing mutations.

N terminal *RPGR* mutations are associated with more severe human disease than those in the C terminal ORF15 domain.(24) However, mouse genetic background has been shown to influence disease severity.(25) To determine if mouse disease correlated with that of human, and was therefore an appropriate model for this work, two novel mutations were generated in the same strain of mice, each replicating human pathogenic mutations (see *Methods* and **Extended Data Fig. 1**). One harboured a frameshift mutation in *Rpgr*’s N-terminus (**Extended Data Fig. 1a**; hereby referred to as *Rpgr*^*Ex3d8*^), while the other harboured a truncation mutation in the C-terminal repetitive domain of the retina-specific isoform (**Extended Data Fig. 1a**; hereby referred to as *Rpgr*^*ORFd5*^).

Fundoscopy revealed retinal degeneration, characterised by punctate lesions at the posterior pole (**Extended Data Figs. 2a and 3a**) whilst fundus blue light autofluorescence showed accumulation of autofluorescent material within the retinas of both models (**Extended Data Figs. 2b and 3b**), a clinical biomarker for photoreceptor disease. Electroretinograms (ERGs) that measure retinal photoreceptor function were recorded as mice aged (**Extended Data Figs. 2f and 4**). The dark-adapted scotopic ERG reflects primarily rod photoreceptor function at lower stimulus intensities, whereas the light-adapted photopic ERG reflects cone photoreceptor function. Both models developed significant loss of their scotopic ERG a-wave amplitude (representing rod photoreceptor dysfunction) at 18 months (**Extended Data Figs. 2f and 4**). Cone photoreceptor function (photopic ERG a-wave amplitude) in the R*Rpgr*^*ORFd5*^ mouse remained comparable to wild type but was significantly reduced from 6 months of age in the *Rpgr*^*Ex3d8*^ mutant (**Extended Data Fig. 4**). Optical coherence tomography showed loss of the outer nuclear (photoreceptor) layer at 18 months (**Extended Data Figs. 2c and 3c**), which was supported by conventional histology in both models but was more severe in *Rpgr*^*Ex3d8*^ (**Extended Data Figs. 2d**,**e and 3d**,**e**). In summary, loss of all Rpgr isoforms leads to more severe disease than loss of the retinal specific isoform alone, similar to human patient mutations. The *Rpgr*^*Ex3d8*^ mouse was brought forward for further studies.

We have previously shown *RPGR* mutations perturb actin regulation in both mouse and induced pluripotent stem cell-derived human photoreceptors.(15) Further, *RPGR* mutations result in loss of the retinal ellipsoid and interdigitating zones of XLRP patients on optical coherence tomography imaging (**Extended Data Figs. 5**); signifying OS loss. We therefore assessed the outer segments of our *Rpgr*-mutant mouse lines using transmission electron microscopy (TEM). At 6 weeks of age, long before the mice show signs of retinal stress **Extended Data Figs. 2g**) and over 12 months before they undergo retinal degeneration (**Extended Data Figs. 2d**,**e and 3d**,**e**), mutant outer segments display a ‘split disc’ phenotype. Their discs have lost compaction and appear spaced out (**Fig. 2a**). Further, mutant outer segments are significantly shorter from base to RPE (**Fig. 2b**) than wild type, suggesting *Rpgr*^*Ex3d8*^ outer segments contain fewer discs. This conclusion is supported by proteomic data of 6 week mouse isolated rod outer segments and 3 month isolated retinas, which show *Rpgr*^*Ex3d8*^ retinas contain lower amounts of photoreceptor disc component (PRCD, Progressive Cone-Rod Dystrophy) (**Fig. 2d, Supplementary Tables. 1 and 2**), a 6 kDa membrane protein that localises exclusively to photoreceptor discs and segregates to the outer disc rim.(26–28) Reduced PRCD in *Rpgr*^*Ex3d8*^ retinas was confirmed on immunoblotting (**Fig. 2e,f**). Basal discs at the base of the *Rpgr*^*Ex3d8*^ outer segment, however, are able to form (**Fig. 2c**).

**Fig. 2.**
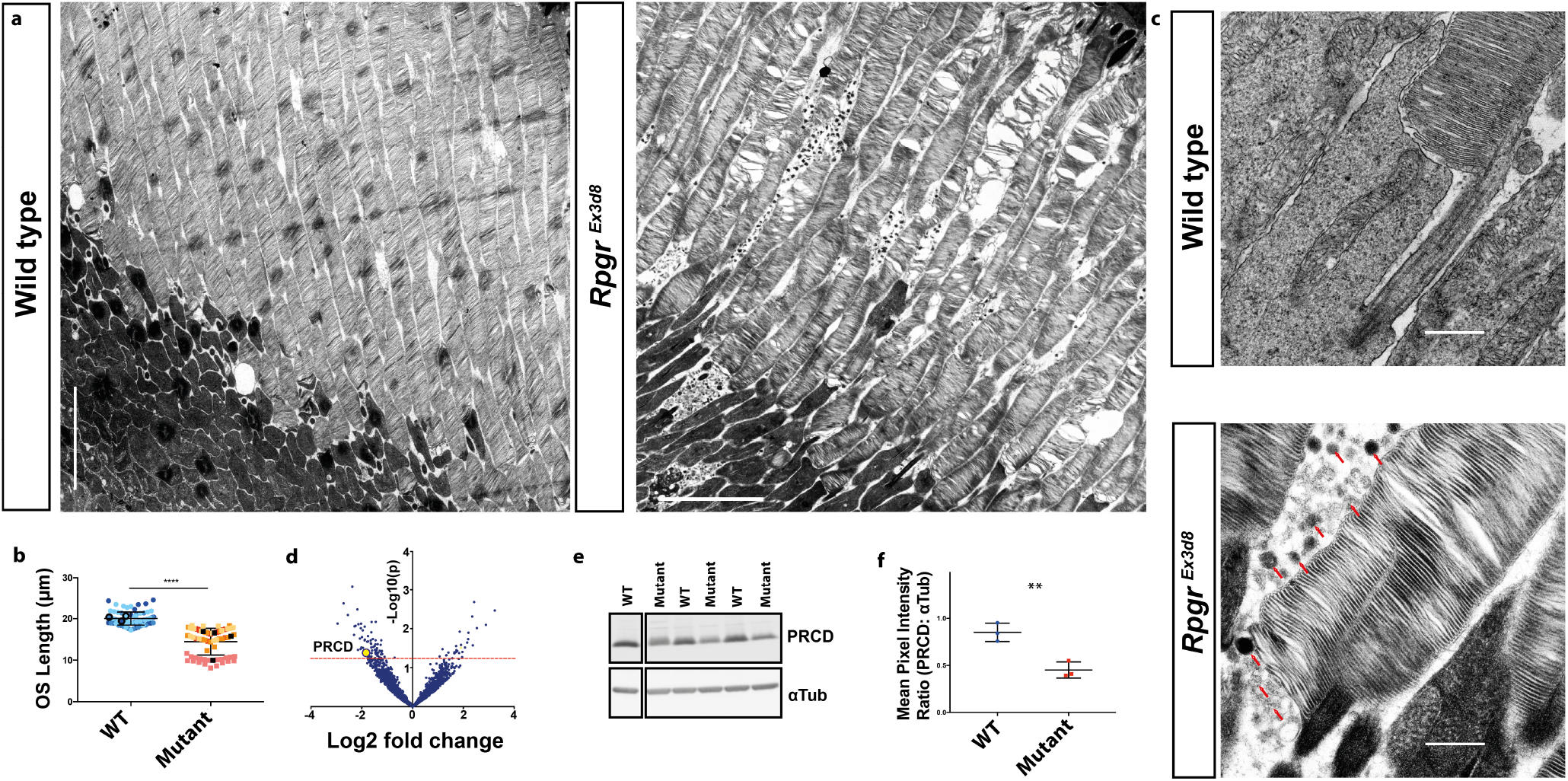
Mutations in *Rpgr* lead to early structural compromise of photoreceptor outer segments (OS). **(a)** Transmission electron micrograph of 6 week old wild type photoreceptors show OS composed of compacted discs extending to the underlying retinal pigment epithelium (RPE) (left panel). Disc compaction is compromised in age-matched mutant photoreceptors, with discs appearing split and spaced out (right panel). **(b)** OS are shorter in *Rpgr*^*Ex3d8*^ mice (black circles denote mean measurement of OS length in each WT experimental animal; black squares denote mean measurement of OS length in each mutant experimental animal; n = 3 animals per genotype; **** = p<0.0001).**(c)** Examination of basal discs at OS base shows compacted discs but an accumulation of shed vesicles (red arrows) in *Rpgr*^*Ex3d8*^ mice. **(d)** Mass spectrometry comparing protein composition of wild type versus *Rpgr*^*Ex3d8*^ 3 month old retinas shows reduced expression of the outer segment protein PRCD in mutant mice. Red line denotes cut off p value for significance. **(e)** Immunoblotting of whole retina lysates confirms reduced PRCD in mutant mice, in keeping with reduced outer segment lengths. **(f)** Quantification of intensity of PRCD relative to loading control alpha tubulin; n=3 per group; ** = p<0.005). (Scale bars; a = 5 *µ*m; c = 0.5 *µ*m).

While basal discs are still able to form, a reduced rate of disc formation could account for the split disc phenotype. *Rpgr*^*Ex3d8*^ mutants accumulated high levels of vesicles in the extra cellular space at the base of outer segments (**Fig. 2c**). Vesicles of similar morphology were recently determined to be ciliary ectosomes in a RDS/peripherin mutant mouse (*rds*^*-/-*^), which displays defective disc formation.(13) Abnormal ciliary membrane deformation, therefore, with compromised discs instead shed as ectosomes, could slow the rate of completed disc formation and account for this split disc phenotype. RPGR is located at the site of disc formation in the distal CC (**Fig. 3a**) and could therefore play a role in disc genesis. To test this, we engineered a novel reporter mouse that would allow tracking of outer segment turnover. The C terminal end of the photosensitive GPCR, Rhodopsin, was tagged with the self-labelling peptide SNAP using CRISPR/Cas9 genome editing to target single cell embryos (**Extended Data Figs. 6a**,**b**). These mice will hereinafter be referred to as *Rhod*^*SNAP*^. Rhodopsin is the major protein component of outer segment discs, and so incubating *Rhod*^*SNAP*^ retinal slice cultures with SNAP fluorophores results in outer segment labelling (**Extended Data Figs. 6c**). The heterozygous SNAP tag knock-in is non-disruptive; heterozygous *Rhod*^*SNAP*^ mice have normal retinal function at 6 months (**Extended Data Figs. 6d**,**e**). Temporal *Rhod*^*SNAP*^ labelling with different SNAP tags *in vivo* allowed optimisation of a block-chase experiment to determine the rate of disc formation over 72 hours (**Fig. 3b, Extended Data Fig. 6f**,**g**) and so were crossed with the *Rpgr*^*Ex3d8*^ mouse. At 6 weeks of age, *Rpgr*^*Ex3d8*^ mice had a reduced rate of new disc formation (**Fig. 3c-e**). We conclude that perturbation of RPGR results in a slowed rate of disc formation, leading to shortened, ‘split disc’ outer segments and vesicle shedding.

**Fig. 3.**
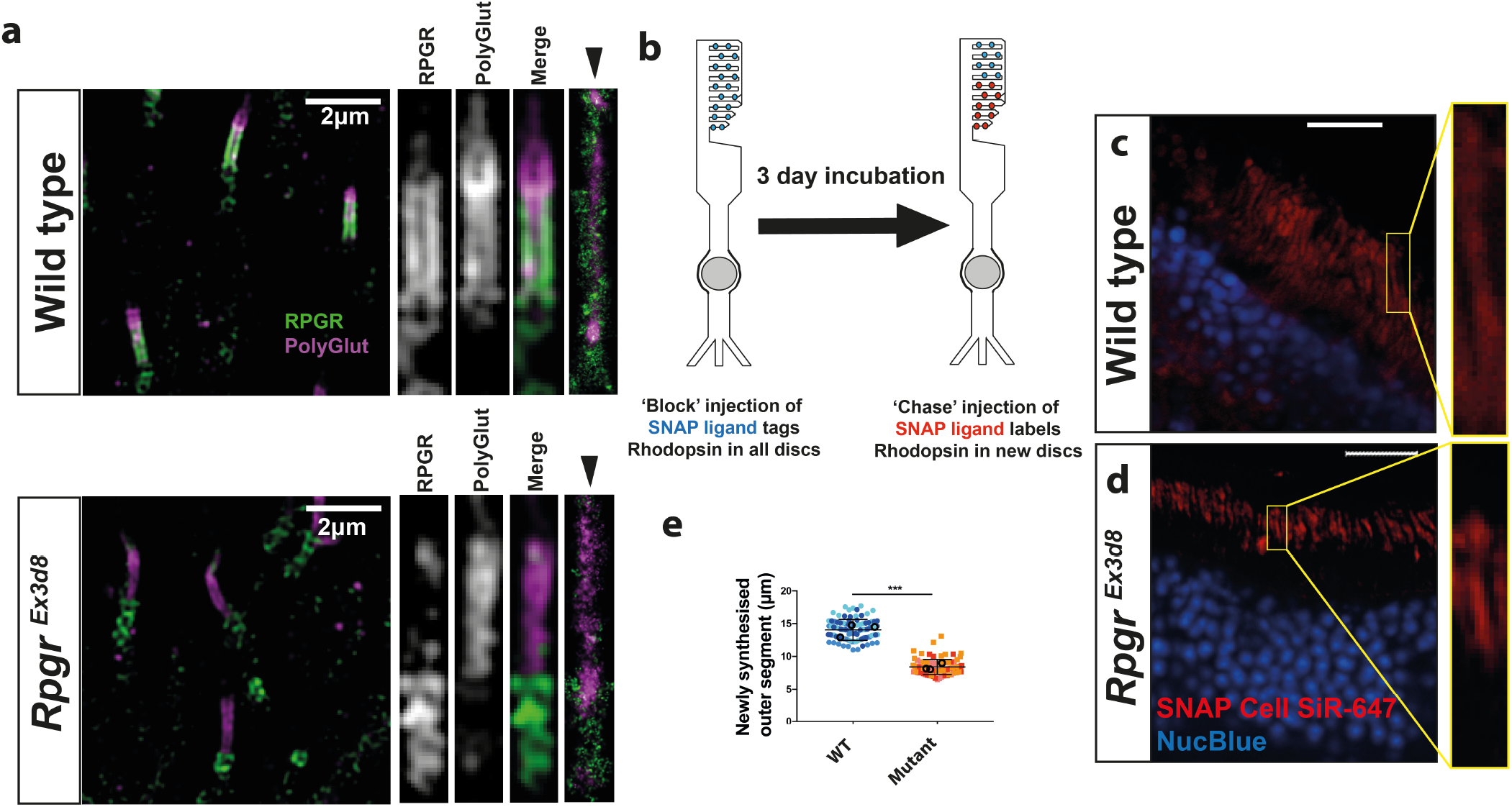
Loss of *Rpgr* reduces the rate of photoreceptor disc formation. **(a)** Top panel: SIM images of wild type photoreceptors show localisation of RPGR’s retinal-specific isoform throughout the length of the photoreceptor connecting cilium, as evidenced by co-localisation with polyglutamylation staining. STORM images (rightmost panel, arrowhead) show RPGR localisation at the ciliary membrane. Bottom panel: Examination of *Rpgr*^*Ex3d8*^ photoreceptors show loss of RPGR staining at the connecting cilium. (NB. green staining below polyglutamylation labelling represents non-specific centrosomal staining). **(b)** Creation of a *Rhod*^*SNAP*^ knock in mouse allows measurement of rate of disc formation. Intravitreal delivery of blocking agent irreversibly binds all Rhodopsin-SNAP molecules. Sequential intravitreal injection of fluorescent SNAP ligand allows labelling of newly synthesised Rhodopsin-SNAP molecules, and therefore new discs. **(c**,**d)** Fluorescent imaging after block-chase procedure shows staining of newly made discs with fluorescent SNAP ligand, 3 days after initial block in wild type (c) and mutant (d) retinas. **(e)** Measurements of block-chase experiment, measuring ‘chased’ labelling of newly formed outer segments, shows a slowed rate of disc formation when *Rpgr* is perturbed. (Scale bars; a = 2 *µ*m; c,d = 20 *µ*m)

### *Rpgr* mutations lead to dysregulation of cofilin activity

Previous work by ourselves and others supports a role for RPGR in regulating photoreceptor actin.(15, 21, 23) To further explore this, we undertook an unbiased total proteomic approach in control and *Rpgr* mutant retinal extracts (**Fig. 4a**). At 3 months of age, computational analysis of differentially expressed proteins using Enrichr(29–31) ranked retinitis pigmentosa as the top disease in *Rpgr*^*Ex3d8*^ retina compared to wild type control (**Extended Data Fig. 7a**). Further, analysis of protein pathways using Panther ranked cytoskeletal regulation by Rho GTPases as a significantly disrupted pathway (**Extended Data Fig. 7a, Supplementary Tables. 1 and 2**). Of note, there was altered expression of the ARP2/3 complex protein ARP3, known to play a role in disc morphogenesis.(10) To further explore this, immunoblotting was carried out on separate total retinal extracts, probing for a panel of actin nucleators and polymerisers. No changes were seen in levels of Cdc42, profilin, Vasp or Wave proteins in *Rpgr*^*Ex3d8*^ retinas (**Extended Data Fig. 7b**), but there was a small, significant reduction in the ARP2/3 complex protein ARP2 (**Fig. 4b**).

**Fig. 4.**
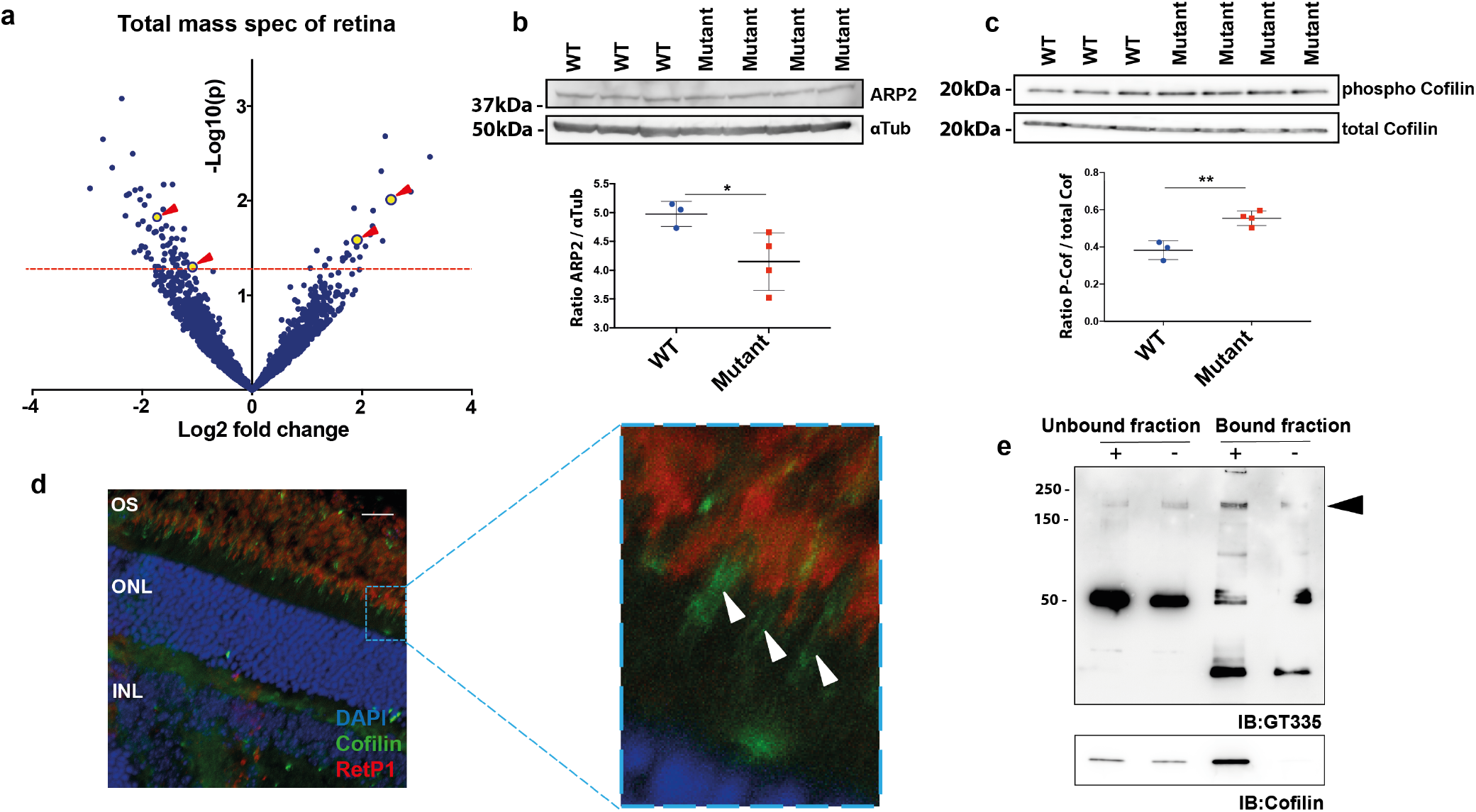
RPGR binds and regulates activity of the actin severing protein cofilin. **(a)** Mass spectrometry analysis of *Rpgr*^*Ex3d8*^ retina shows dysregulation of actin binding proteins (labelled in yellow; red arrows). **(b)** Immunoblotting shows modest reduction of ARP2 in *Rpgr*^*Ex3d8*^ retina lysates (y axis denotes ratio of ARP2 to *α*-tubulin loading control). **(c)** Immunoblotting shows increased phosphorylation of cofilin at Serine 3 (and therefore reduced activity) (y axis denotes ratio of phospho-cofilin to total cofilin). **(d)** Immunohistochemistry of an OS marker, RetP1, and cofilin show cofilin localisation to the photoreceptor connecting cilium (white arrowheads) in wild type retinas. **(e)** Immunoprecipitation of wild type retinal lysates using magnetic beads coated (+) or uncoated (-) with cofilin antibody shows enrichment of cofilin in bound fraction (bottom panel) and the retinal specific isoform of RPGR, as detected using GT335 antibody (black arrowhead, top panel; band at 52 kDa is acetylated tubulin). (Scale bar; d = 20 *µ*m)

The ARP2/3 complex directly competes for binding sites on actin filaments with the actin depolymeriser cofilin, which we have previously shown to have reduced activity in patient-derived *RPGR*-mutant, iPSC retinal organoid cultures(12) and is thought to be present in early discs.(10) To this end, we analysed *Rpgr*^*Ex3d8*^ retinal lysates with immunoblotting, which showed increased phosphorylation, and therefore inactivation, of cofilin at Serine 3 (**Fig. 4c**). Using immunofluorescence, we demonstrate that cofilin localises to the photoreceptor CC membrane (**Fig. 4d**), at the site of disc morphogenesis and overlapping with RPGR localization (**Fig. 3a**). We therefore sought to determine whether there is a biochemical interaction between RPGR and cofilin in vivo. Endogenous co-immunoprecipitation (co-IP) experiments using murine retinal lysates confirmed an interaction between endogenous cofilin and the retinal specific isoform of RPGR (**Fig. 4e**).(32) Given actin’s multifunctionality and its abundance throughout the cell, it is crucial that local, sub-cellular turnover is tightly controlled to ensure cellular processes are precisely executed, both temporally and spatially. Our findings support that RPGR binds and regulates actin binding protein activity in the distal CC to regulate basal disc formation.

### Perturbation of RPGR dysregulates actin-mediated photoreceptor disc formation

Having demonstrated dysregulation of actin binding proteins in *Rpgr*^*Ex3d8*^ mice, and a reduced rate of disc formation, we examined local actin dynamics in the CC in the absence of RPGR. We confirmed increased filamentous actin was seen at the base of the OS in *Rpgr*^*Ex3d8*^ compared to wild type (**Extended Data Fig. 7c-e**).(12) To better resolve actin bundle integrity at the site of basal disc formation, *Rpgr*^*Ex3d8*^ photoreceptors were further examined using cryoET. At 3 months of age, in keeping with reduced severing activity of cofilin, analysis revealed an increase in the length of actin microfilaments in *Rpgr*^*Ex3d8*^ mice compared to wild type (**Fig. 5a-c**). However, cryoET provides only a snapshot of the very dynamic process of actin turnover. To this end, we examined the rate of change of the actin microfilaments in *Rpgr*^*Ex3d8*^ and wild type mice as captured by live imaging of retinal slice cultures using a live actin label, SiR-Actin. Analysis showed reduced actin dynamics in the CC of *Rpgr*^*Ex3d8*^ photoreceptors (**Fig 5d**,**e, Extended Data Movies. 1 and 2**).

**Fig. 5.**
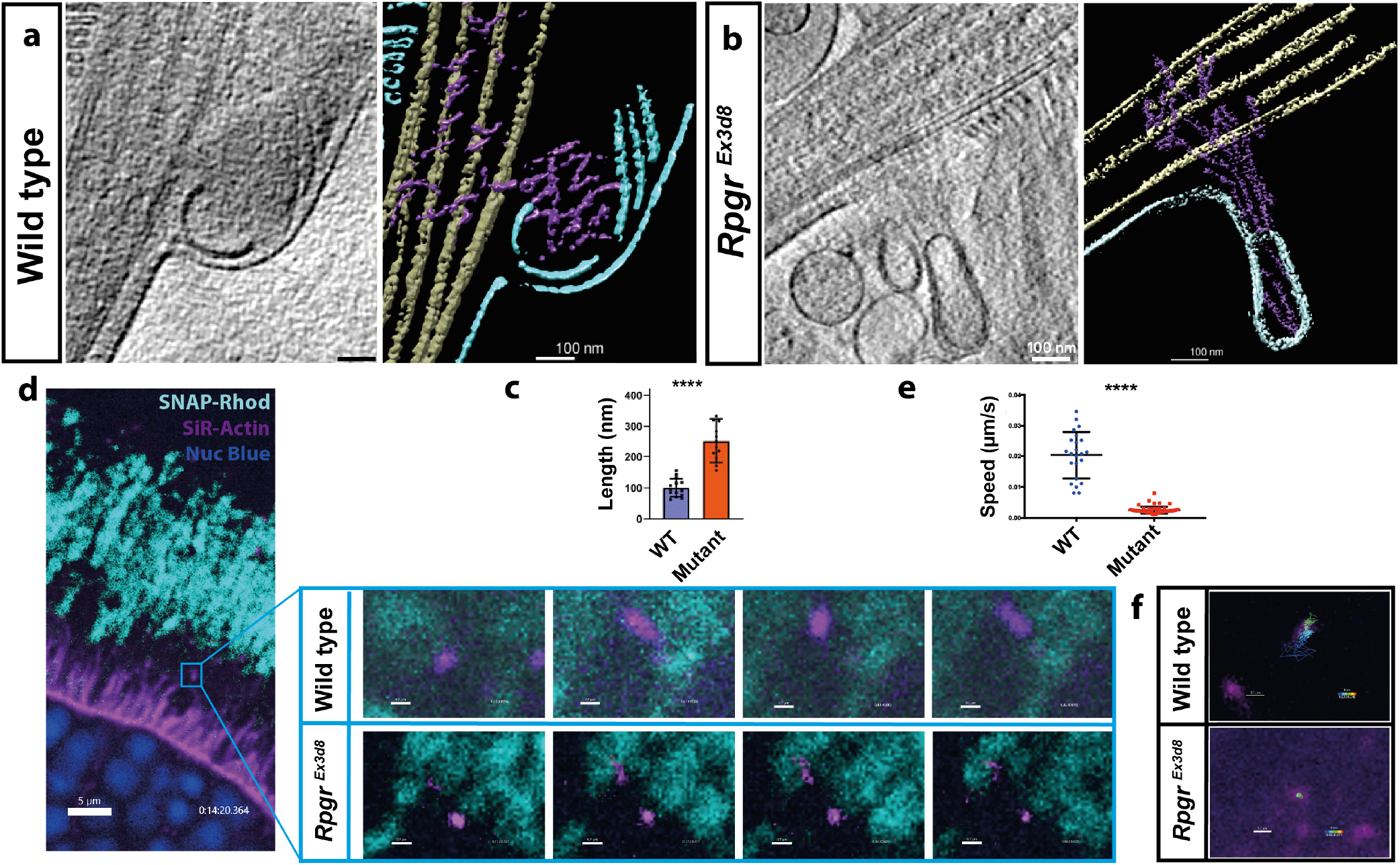
Loss of Rpgr hyperstabilises actin in the basal photoreceptor disc and reduces actin dynamics. **(a**,**b)** A projection from a tomogram (left panel) and a slice through a tomographic reconstruction (right panel) of the basal disc of a wild type (a) and *Rpgr*^*Ex3d8*^ (b) photoreceptor to highlight the ciliary and disc membranes (cyan), microtubule based axoneme (tan) and actin filaments (purple). **(c)** Measurement of actin filament length in the basal disc of photoreceptor tomograms shows them to be increased in the *Rpgr*^*Ex3d8*^ mouse (n = 4 per group; **** = <0.0001). **(d)** Live imaging of retinal slice cultures using the *Rhod*^*SNAP*^ mouse allows visualisation of nuclei (blue) and connecting cilium actin bundles (magenta) at the base of the outer segments (cyan) over time. Sample images at high magnification show actin bundle dynamics in wild type (top panel) and *Rpgr*^*Ex3d8*^ (bottom panel) photoreceptors. **(e)** Tracking of actin bundle movement by live cell imaging in culture shows reduced dynamics in *Rpgr*^*Ex3d8*^ retinas (speed = distance moved in x axis / time measured; n = 3 per group; **** = < 0.0001). **(f)** Sample images of tracked connecting cilium actin bundles in wild type (top panel) and *Rpgr*^*Ex3d8*^ (bottom panel) photoreceptors over a 60 minute period. ‘Dragon tail’ projection depicts distance moved in this time. (Scale bars; a,b = 100nm; d (large panel) = 5 *µ*m; d (small panels) = 0.7 *µ*m; f = 0.7 *µ*m)

In the absence of RPGR, we have shown increased levels, steady state lengths and slowed turnover of filamentous actin in the region of the newest basal discs, consistent with the altered actin dynamics leading to some nascent discs being aborted and being shed as the observed vesicles/ectosomes (**Fig. 2c**). We reasoned, therefore, that depolymerising actin in mutant retinas could rescue the disc biogenesis phenotype. *Rpgr*^*Ex3d8*^ mice were treated for 6 hours with the actin depolymeriser, cytochalasin D, by intravitreal injection of 0.5 *µ*l at 25 mM. TEM analysis revealed a significant reduction in the number of vesicles shed at the base of the mutant outer segments with treatment (**Fig. 6a,b**), as well as an overgrowth of the basal discs that did form as previously described(9, 10). With this rescue experiment, we conclude that the shedding of aborted discs as vesicles and a reduced rate of disc formation in *RPGR* mutants is due to compromised dynamic turnover of actin microfilaments extending from the photoreceptor CC into basal discs, due to cofilin dysregulation. Our results suggest that these defects in the cadence of disc biogenesis, resulting from increased length of these mutant actin microfilaments, can be therapeutically modulated short-term.

**Fig. 6.**
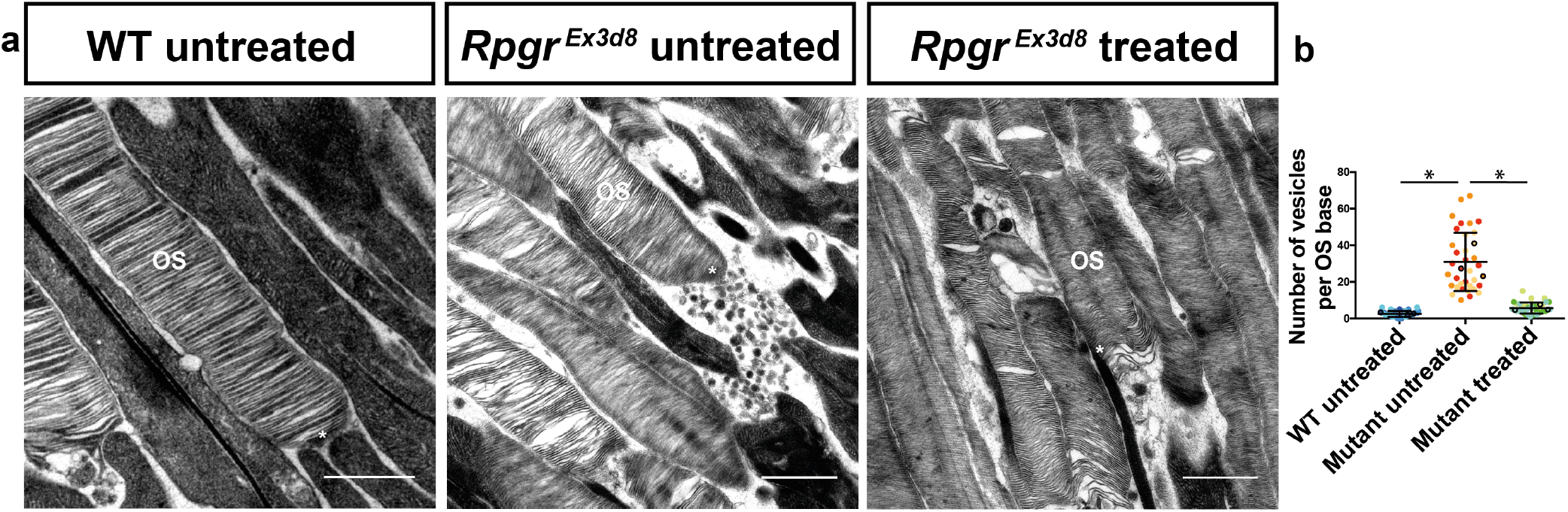
Treatment with an actin depolymerising agent rescues the vesicles shed from Rpgr-mutant photoreceptors. **(a)** The *Rpgr*^*Ex3d8*^ mouse sheds high numbers of vesicles from the base of the outer segment (middle panel) compared to wild type (left panel). Intravitreal delivery of the actin depolymerising agent, cytochalasin D, rescues this phenotype, significantly reducing the number of shed vesicles (right panel)(scale bars = 1 *µ*m; OS = Outer Segment; * = outer segment base).**(b)** Quantification showing significantly increased vesicle shedding at the base of the *Rpgr*^*Ex3d8*^ photoreceptors, which is rescued by cytochalasin treatment. (Black circles denote mean number of vesicles at the base of each photoreceptor per experimental animal; n = 3 animals per genotype).

## Discussion

Our *in vivo* data support that RPGR functions locally at the photoreceptor CC to spatially regulate activity of actin severing proteins, such as cofilin, in order to appropriately pace actin-mediated disc formation. We have previously shown that RPGR interacts with another actin severing protein, gelsolin, and these interactions are perturbed in pathogenic human RPGR retinal organoids(12). We propose, therefore, that the glutamylated RPGR isoform serves as a structural scaffold that facilitates the timely recruitment and activation of actin binding proteins (ABPs) to control the appropriate pace of disc biogenesis (**Extended Data Fig. 8**). Our study raises important questions about what the molecular nature of this pacemaker is that controls the regular tempo. Previous work at the primary cilia tip has demonstrated the highly localised role that actin plays in ciliary membrane deformation in a content-dependent manner. Here, excision of a membranous bud (ectosome) at its distal end, facilitates the exit of GPCRs to regulate cilia content appropriately(1, 2). Similar mechanisms at the distal CC / transition zone may operate to ‘sense’ regulate membrane content, such as rhodopsin concentration, to regulate disc formation. Further studies are necessary to distinguish these models. We propose that photoreceptor disc formation shares a common ancestral mechanism of ectocytosis and RPGR’s retinal specific isoform may have evolved to promote this.

It is over 30 years since disc formation was first proposed to be an actin-dependent process. In seminal TEM experiments, it was shown that actin was present in the distal CC(8) and that depolymerising actin inhibited the formation of new discs, but not the addition of membrane to existing ones,(9) suggesting actin bundling was required for the initial membrane deformation to begin the process. Subsequently, it was confirmed that the process of disc morphogenesis is one of evagination(33–35), which would support a process of active, actin protrusions initiating the process. In recent years, progress has been made in defining the molecular pathways that govern actin regulation in the CC(10, 11, 13, 15). Here, for the first time, we confirm the presence of actin microfilament bundles within the basal disc and, by live imaging, demonstrate the dynamic changes in actin orientation that occur precisely at the level of the distal CC.

Force generation lies at the heart of all cellular membrane morphogenesis, which the actin cytoskeleton can produce due to its ability to form higher order structures. But these resulting structures can remain stable for an incredibly long time.(36) In order for cells to perform the rapid actin turnover critical for a metronomic, repeated membrane deformation such as that required for disc formation, regulated dismantling of these networks is equally important. Thus, it would be predicted that a hyperstabilised actin cytoskeleton at the site of basal disc formation, as seen in our *Rpgr*-mutant mice, would compromise the cadence of disc genesis, which we demonstrate. The actin severing protein, cofilin, plays a central role in actin-filament length oscillation by trimming the aged end of a filament that continues to grow at the barbed end,(36) allowing branched networks to be disassembled.(37) Of interest, cofilin competes for binding to actin with the actin polymeriser, ARP2/3, a protein previously demonstrated to be crucial for disc formation.(10) Cofilin is known to regulate primary cilia length through actin rearrangement.(38) Our work suggests cofilin is required for mammalian photoreceptor disc formation, and the observation that cofilin is present in aborted photoreceptor ectosomes when normal disc formation is disrupted(10) further supports this model.

The molecular architecture of this RPGR-dependent scaffold at the distal CC involves SPATA7 and RPGRIP,(39) mutations in which also cause RP in humans. RPGR is mislocalised away from the distal CC in mice lacking either SPATA7(39) or RPGRIP,(40) with failure of normal outer segment disc genesis. Importantly, while RPGR, RPGRIP and SPATA7 appear critical for photoreceptor function, they appear dispensable for cilia function in most other cell types. We speculate, therefore, that they have evolved as a scaffold at the site of disc formation to which actin binding proteins, critical for timely disc formation, can be recruited. Pharmaco-manipulation of actin activity could reverse the underlying cellular phenotypes disrupting disc biogenesis. Further work is required to address the translational potential of our study, specifically to establish if actin modulation could slow the retinal degeneration seen in *RPGR/XLRP* patients.

## Methods

### Animal experiments

All experiments followed international, national and institutional guidelines for the care and use of animals. Animal experiments were carried out under UK Home Office Project License PPL P1914806F in facilities at the University of Edinburgh (PEL 719 60/2605) and were approved by the University of Edinburgh animal welfare and ethical review body.

### Mouse generation

*Rpgr* mutant mice: Using CRISPR/Cas9 genome editing to target single cell embryos, deletion mutations were introduced into the Rpgr locus of C57Bl6/J mice (**Extended Data Fig. 1**). Ensemble *Rpgr* EN-SMUSG00000031174, chromosome X X:10,175,515 was used for CRISPR design. The CRISPR target sequences (20-nucleotide sequence preceded by Bbs1 ends (Fd CACC / Rv AACA), followed by a protospacer adjacent motif [PAM] of ‘NGG’) were selected using the prediction software (www.crispr.mit.edu) and synthesised by Sigma, annealed and ligated into the pX330-U6-Chimeric-BB-CBh-hSpCas9 plasmid (Addgene 42230). The plasmid was transformed into chemically competent Stbl3 cells, colonies grown, maxi-prepped and correct sequencing confirmed by Sanger sequencing. The repair template oligo for RpgrORFd5 (see below) was synthesised by Integrated DNA Technologies. See **Supplementary Table 3**.**3** for list of guides and repair templates.

An 8 base pair deletion was introduced into exon 3 (c.296del8), resulting in a premature termination codon 68 base pairs downstream being shifted into frame and a predicted null protein (**Extended Data Fig. 1**); this mouse will hereby be referred to as *Rpgr*^*Ex3d8*^. In a separate round of injections, a 5 base pair deletion was introduced at the beginning of the repetitive domain of RPGR’s retinal specific splice variant (c.2478del5), resulting in a premature termination codon 52 base pairs downstream being shifted into frame (**Extended Data Fig. 1**) and a predicted truncated protein lacking all of the protein’s glutamate rich consensus motifs (GEEEG), whose glutamylation is crucial for protein function(32). This mouse will hereby be referred to as *Rpgr*^*ORFd5*^.

Genome editing was performed by microinjection of the *Rpgr* guide / Cas9 plasmid (5 ng/*µ*l) with or without a single stranded repair oligo mRpgr (100 ng/*µ*l) into single cells embryos. The injected zygotes were cultured overnight in KSOM for subsequent transfer to the oviduct of pseudopregnant recipient females.(41) From microinjection CRISPR targeting, genomic DNA from founder ear-clip tissue was subject to PCR screening and subsequent sequence verification of successful mutagenesis events by Sanger sequencing (**Extended Data Fig. 1**). Founder mice, mosaic for desired mutations were crossed with CD1 mice to generate large numbers of F1 progeny and allow unwanted allelic variants to segregate. Selected F1 mutants were backcrossed onto C57 mice and subsequently intercrossed. Initial genotyping was performed by PCR and Sanger sequencing using primers outlined in **Supplementary Table 3**.**3**. The genotyping of subsequent animals was performed by TransNetyx.

Western blot analysis showed loss of detection of a polyglutamylated protein running at the weight of Rpgr’s retinal isoform (**Extended Data Fig. 1**)(32). In the absence of a commercially available antibody for murine RPGR, we consider this as confirmation of loss of protein expression.

*Rhodopsin-SNAP (Rhod*^*SNAP*^) mouse: Using CRISPR/Cas9 genome editing to target single cell embryos, a SNAP tag was introduced at the C terminal end of the Rhodopsin locus of C57Bl6/J mice (**Extended Data Fig. 6**). Ensemble *Rhodopsin* ENSMUST00000032471.9, Chromosome 6: 115,908,709-115,916,997 was used for CRISPR guide design, selected using the prediction software (www.crispr.mit.edu) and synthetic crRNA synthesised by Dharmacon for use with their Edit-R tracrRNA technology. A repair plasmid, with SNAP targeted to Rhodopsin’s C terminus, was synthesised by GeneArt (Invitrogen). crRNA and tracrRNA was annealed and injected with the repair plasmid and Cas9 protein (Invitrogen 825641 Geneart Platinum). See **Supplementary Table 3**.**3** for list of guides and repair templates.

Genome editing was performed by microinjection of the *Rhodopsin* guide/ Cas9 protein/repair plasmid mix into single cells embryos. The injected zygotes were cultured overnight in KSOM for subsequent transfer to the oviduct of pseudopregnant recipient females.(41) From microinjection CRISPR targeting, genomic DNA from founder ear-clip tissue was subject to PCR screening and subsequent sequence verification of successful mutagenesis events by Sanger sequencing (**Extended Data Fig. 6**). Founder mice, mosaic for desired mutations were crossed with CD1 mice to generate large numbers of F1 progeny and allow unwanted allelic variants to segregate. Selected F1 mutants were backcrossed onto C57 mice and subsequently intercrossed. Initial genotyping was performed by PCR and Sanger sequencing using primers outlined in **Supplementary Table 3**.**3**. The genotyping of subsequent animals was performed by TransNetyx.

Successful SNAP-mediated dye incorporation in the knock in was confirmed by labelling rhodopsin in the photoreceptor outer segments with fluorescent SNAP probes (**Extended Data Fig. 6**).

### Electrodiagnostic testing

All mice undergoing EDT were dark adapted overnight prior to the procedure, and experiments were carried out in a darkened room under red light using an HMsERG system (Ocuscience). Mice were anesthetised using isofluorane and pupils were dilated through the topical application of 1% w/v tropicamide before being placed on a heated ERG plate. Three grounding electrodes were used subcutaneously (tail, and each cheek) and silver embedded electrodes were placed upon the cornea held in place with a contact lens. The standard International Society for Clinical Electrophysiology of Vision (ISCEV) protocol was used, which recorded scotopic responses before a 10 min light adaptation phase in order to record photopic responses(42). 3 and 10 cd.s/m^−2^ light intensity scotopic responses were used for analysis. Data was analysed using Graphpad Prism and compared by unpaired t-test.

### Optical coherence tomography (OCT) and blue light autofluorescence imaging

Mice were anesthetised using isofluorane and pupils were dilated through the topical application of 1% w/v tropicamide before being placed on a custom built, heated plate on a Heidelberg Spectralis OCT machine. Mouse retina was brought into focus using IR imaging and fast volume scans (20° x 20°) obtained (25 sections; 240 *µ*m between sections) by spectral domain OCT using a 55 dioptre lens adapted for small animals. Accumulation of autofluorescent material in the retina, a clinical biomarker for retinal dysfunction, was assessed using the BluePeak™ blue light function on the Spectralis.

### Protein extraction, antibodies and western blotting

Retinas were lysed in 50 mM Tris pH8.0, 150 mM NaCl, 1% NP-40 buffer containing protease inhibitors. Protein samples were separated by SDS-PAGE and electroblotted onto nitrocellulose membranes (Whatman) using the iblot System for 6 min (Invitrogen). Non-specific binding sites were blocked by incubation of the membrane with 5% non-fat milk in TBS containing 0.1% Tween 20 (TBST) for 1 hour. Proteins were detected using the primary antibodies diluted in blocking solution (4% Bovine Serum Albumin in TBST; **Supplementary Table 3**.**1 and 3**.**2** for list of antibodies). Following washing in PBST, blots were incubated with the appropriate secondary antibodies conjugated to horse-radish peroxidase (Pierce) and chemiluminescence detection of Super Signal West Pico detection reagent (Pierce) by high resolution image capture using the ImageQuant LAS4000 camera system (GE Healthcare). Images were transferred to Fiji/ImageJ and mean pixel intensity of protein bands measured for quantification, with an equal area of blot assessed across all bands.

### Rod outer segment proteomics

6 week old, dark-adapted mice were maintained in constant darkness for 12 hr overnight prior to retina harvesting (also performed in the dark with infrared illumination). Rod outer segments were isolated by vortexing retinas in 150ul of 27% sucrose solution in 20 mM HEPES, pH 7.4, 100 mM KCl, 2 mM MgCl2 and 0.1 mM EDTA, spinning at 200 g for 30 sec and collecting supernatant. Supernatant was diluted with the same buffer without sucrose and loaded in a step gradient of 27% and 32% sucrose prior to spinning at 25,000 rpm and outer segment collection from the 32-27% sucrose interface. Proteins were solubilized in 2% SDS, 100 mM Tris·HCl (pH 8.0), reduced with 10 mM DTT (D0632; Sigma-Aldrich), alkylated with 25 mM iodoacetamide (I1149; Sigma-Aldrich), and subjected to tryptic hydrolysis using the HILIC beads SP3 protocol(10). The resulting peptides were analyzed with a nanoAcquity UPLC system (Waters) coupled to an Orbitrap Q Exactive HF mass spectrometer (Thermo Fisher Scientific) employing the LC-MS/MS protocol in a data-independent acquisition mode. The peptides were separated on a 75-µm × 150-mm, 1.7-*µ*m C18 BEH column (Waters) using a 120-min gradient of 8 to 32% of acetonitrile in 0.1% formic acid at a flow rate of 0.3 mL/min at 45 °C. Eluting peptides were sprayed into the ion source of the Orbitrap Q Exactive HF at a voltage of 2.0 kV. Progenesis QI Proteomics software (Waters) was used to assign peptides to the features and generate searchable files, which were submitted to Mascot (version 2.5) for peptide identification. For peptide identification, we searched against the UniProt reviewed mouse database (September 2019 release) using carbamidomethyl at Cys as a fixed modification and Met oxidation as a variable modification. Proteins were included based on 3 criteria: 1) at least 1 peptide is identified in each independent experiment, 2) at least 2 peptides are identified in one experiment, and 3) the identification confidence score is >95% in both experiments.

### Whole retina proteomics

3 month old, dark-adapted mice were maintained in constant darkness for 12 hr overnight prior to retina harvesting (also performed in the dark with infrared illumination). Retinas were lysed in 2% SDS and kept at -80 °C until testing. 2 retinas from 1 mouse were used per biological replicate, with 5 biological replicates per group (mutant and wild type). In all cases, cell lysates were digested using sequential digestion of LysC (Wako) and trypsin (Pierce) using the FASP protocol (https://pubmed.ncbi.nlm.nih.gov/25063446/). Samples were acidified to 1% TFA final volume and clarified by spinning on a benchtop centrifuge (15k g, 5 min). Sample clean-up to remove salts was performed using C18 stage-tips (Rappsilber et al., 2003). Samples were eluted in 25 µl of 80% acetonitrile containing 0.1% TFA and dried using a SpeedVac system at 30 °C and resuspended in 0.1% (v/v) TFA such that each sample contained 0.2 *µ*g/ml. All samples were run on an Orbitrap FusionTM Lumos mass spectrometer coupled to an Ultimate 3000, RSL-Nano *µ*HPLC (both Thermo Fisher). 5 *µ*l of the samples were injected onto an Aurora column (25 cm, 75 *µ*m ID Ionoptiks, Australia) and heated to 50 °C. Peptides were separated by a 150 min gradient from 5-40% acetonitrile in 0.5% acetic acid. Data was acquired as data-dependent acquisition with the following settings: MS resolution 240k, cycle time 1 s, MS/MS HCD ion-trap rapid acquisition, injection time 28 ms. The data were analyzed using the MaxQuant 1.6 software suite (https://www.maxquant.org/) by searching against the murine Uniprot database with the standard settings enabling LFQ determination and matching. The data were further analyzed using the Perseus software suite. LFQ values were normalized, 0-values were imputed using a normal distribution using the standard settings.

### Immunohistochemistry

Mice were sacrificed and eyes enucleated and placed into Davidson’s fixative (28.5% ethanol, 2.2% neutral buffered formalin, 11% glacial acetic acid) for 1 hour (cryosectioning) or overnight (wax embedding). For cryosectioning, eyes were removed from Davidson’s fixative and placed into a series of consecutive 10%, 15% and 20% sucrose in PBS buffer for 15 mins, 15 mins and overnight respectively to cryopreservation. Eyes were then embedded using OCT and kept at -80 °C until sectioned. Sections/cells were blocked/permeabilised with 4% BSA (Sigma) and 0.5% Triton X100 (Fisher) for 1 hour at RT, washed with PBS, incubated with primary antibodies overnight at 4 °C, washed in PBS, incubated with secondary antibodies for 60 min at RT, washed in PBS, incubated in Hoechst for 5 min at RT and mounted with coverslips using Fluoromount-G (Southern Biotech). For wax preservation, eyes were removed from Davidson’s fix and dehydrated through an ethanol series of 70% ethanol, twice in 70%, the 80% xyelene, thebn 90% paraffin and finally twice in 100%; paraffin each for 45 mins. Hematoxylin and eosin staining was performed on 8 µm paraffin tissue sections and imaged on a Zeiss Brightfield microscope. Confocal imaging was performed on a Nikon Eclipse TiE inverted microscope with Perfect Focus System using resonant scanning mirrors (equipped with 405nm diode, 457/488/514nm Multiline Argon, 561nm DPSS and 638nm diode lasers) with detection via four Photomultiplier tubes (2x standard Photomultiplier tubes and 2x GaAsP PMTs). A 60x oil immersion lens was used. Data was acquired using NIS Elements AR software (Nikon Instruments Europe, Netherlands). Z-stacks were processed and analysis in Fiji (ImageJ).

### Airyscan imaging

Sample preparation for Airyscan confocal immunofluorescence microscopy. Mice 6 weeks of age were transcardially perfused with 80 mM PIPES, pH 6.8, 5 mM EGTA, 2 mM MgCl2, 4% paraformaldehyde). Eyes were enucleated, and after the cornea was removed, they were immersion fixed overnight at 4 °C. After removal of lens, the eyecups were flash-frozen in optimal cutting temperature (OCT) using liquid nitrogen. 8 *µ*m cryosections were collected and stained for rhodamine wheat germ agglutinin (Vector RL-1022) and phalloidin conjugated to Atto647N (Sigma 65906). The sections were then mounted in Prolong Glass (Invitrogen P36980). Sections were imaged on a Zeiss LSM 880 Airyscan Fast Confocal Microscope using a 63x objective. Z-stacks were first processed in Zeiss ZenBlue software for Airyscan processing, then colour processing and analysis were performed in Fiji/ImageJ. Actin puncta quantification was performed by taking a 0.35 *µ*m ROI around each actin puncta that was at the base of an outer segment, slice by slice, on each z-stack. Then the averaged relative integrated density of three background ROI’s were subtracted from each relative integrated density measurements from the actin ROIs. These measurements were then plotted in Prism software, where Students t-test statistical analysis was performed.

### Transmission electron microscopy

6 week old mice were euthanized by transcardial perfusion using fixative (50mM MOPS, pH 7.4, 2% glutaraldehyde, 2% paraformaldehyde, 2.2 mM CaCl2). Eyecups were enucleated and placed in 1 mL of fixative. After 30 minutes, the cornea and lens were removed, then left to incubate in fixative for a total of 2 hours at room temperature. Thereafter, retinas were either: 1.Washed in 0.1 M phosphate buffer (pH 7.4), postfixed with 1% osmium tetroxide (ElectronMicroscopy Science) and dehydrated in an ethanol series prior to embedding in Medium Epoxy Resin(TAAB). Ultrathin (75 nm) sections of the retina were then stained with aqueous uranyl-acetate andlead citrate and then examined with a Hitachi 7000 electron microscope (Electron Microscoperesearch services, Newcastle University Medical School).

2. Polymerized in 4% agarose (Genemate E-3126-25). 150 *µ*m sections were collected into MilliQ waterusing a vibratome. Sections were then stained in 1% tannic acid in 0.1 M HEPES, pH 7.4, for 1 hour,with rocking at room temperature, covered. After rinsing in MilliQ water, the sections were stainedwith 1% uranyl acetate in 0.2 M maleate buffer, pH 6.0, for 1 hour, with rocking at room temperature,covered. The sections were rinsed in MilliQ water and dehydrated in a series of ethanol washes (50%, 70%, 90%, 100%, 100%)for 15 minutes each, followed by two 100% acetone washes, 15 minutes each.The sections were then embedded in Epon-12 resin by sandwiching the sections between two sheetsof ACLAR (EMS 50425-10) and leaving them at 60 °C for 48 hours. Ultrathin silver sections (60 nm)were placed on copper slot grids (EMS FF2010-CU) and poststained in 1.2% uranyl acetate in MilliQwater for 6 minutes, followed by staining in Sato’s lead (a solution of 1% lead acetate, 1% lead nitrate,and 1% lead citrate; all from Electron Microscopy Sciences) for 2 minutes. Grids were imaged on aJEOL JEM-1400 electron microscope.

### Cytochalasin rescue experiment

Mice were anesthetised using isofluorane and pupils were dilated through the topical application of 1% w/v tropicamide. 0.5 *µ*l of 25mM Cytochalasin D (Sigma) in PBS was injected intravitreally. Contralateral eyes had a sham injection of 0.5 *µ*m PBS and served as the control eye. 6 hours later mice underwent transcardial perfusion using fixative and processed for electron microscopy as outlined above.

### STORM immunohistochemistry and resin embedding

Unfixed retinas from 6- to 12-week-old wild type mice were immunolabeled for STORM using a protocol previously developed.(43) Most 1 *µ*m resin sections were collected into 35 mm glass-bottom dishes, and 2 sections for each sample were also collected onto a 1.5 coverslip, mounted in Prolong Glass (Invitrogen P36980) and imaged on a Structured Illumination Microscope (SIM); DeltaVision OMX Blaze v4 (GE Healthcare, now Cytiva); a PLANPON6 60×/NA 1.42 (Olympus) using oil with a refractive index of 1.520. Z-spacing of 125 nm was used for each 1 *µ*m z-stack. SIM reconstructions and alignment were performed in Softworx 7 software. After analysis, reconstructions were processed in Fiji/ImageJ.

Immediately prior to STORM imaging, 10% sodium hydroxide (w/v) was mixed with pure 200-proof ethanol for 45 minutes to prepare a mild sodium ethoxide solution. Glass-bottom dishes with ultrathin retina sections were immersed for 30–45 minutes for chemical etching of the resin. Etched sections were then washed and dried on a 50 °C heat block. The following STORM imaging buffer was prepared: 45 mM Tris (pH 8.0), 9 mM NaCl, and oxygen scavenging system: 10mM Sodium Sulfite, 10% (w/v) dextrose + 100 mM MEA (i.e., L-cysteamine, Chem-Impex) + 10% VECTASHIELD (Vector Laboratories). Imaging buffer was added onto the dried, etched sections and sealed with a second #1.5 coverslip for imaging. Imaging was performed on the Nikon N-STORM system, which features a CFI Apo TIRF 100× oil objective (NA1.49) on an inverted Nikon Ti Eclipse microscope. Photobleaching and photoswitching initiation were performed using both the 561 and 647 mm laser lines at maximum power. Imaging frames were collected at 30 frames per second. A total of 40,000 frames were collected for each imaging experiment.

### STORM image analysis

Two-dimensional (2D) STORM analysis of STORM acquisition frames was performed using NIS-Elements Ar Analysis software. Analysis identification settings were used for detection of the individual point spread function (PSF) of photoswitching events in frames from both channels to be accepted and reconstructed as 2D Gaussian data points. These settings were as follows: minimum PSF height: 400, maximum PSF height: 65,636, minimum PSF width: 200 nm, maximum PSF width: 700 nm, initial fit width: 350 nm, maximum axial ratio: 2.5, maximum displacement: 1 pixel. After analysis, reconstructions of single cilia were processed in Fiji/ImageJ for straightening.

### Block chase experiment

Mice underwent intravitreal ‘block’ injection of 1 *µ*l of 0.6 *µ*M SNAP-Cell TMR-Star (New England Biolabs (S9105S)) under inhalational anaesthesia. 72 hours later, mice were sacrificed. Eyes were immediately enucleated and underwent keratectomy, sclerectomy and lensectomy. Retinal cups were embedded in 5% agar moulds and immediately underwent sectioning on a fully automated vibratome (Leica VT1200 S). Sections were ‘Chased’ by incubation with 0.6 *µ*M SNAP-Cell 647-SiR (New England Biolabs (S9102S) in retinal culture media for 90 minutes at 37 °C / 5% CO2. 30 minutes before imaging, NucBlue Live ReadyProbe Reagent (Hoechst 33342; ThermoFisher R37605) was added to the culture for nuclear staining. Imaging of slices was acquired using a x40 oil immersion lens on the multimodal Imaging Platform Dragonfly (Andor technologies, Belfast UK) equipped with 405, 445, 488, 514, 561, 640 and 680nm lasers built on a Nikon Eclipse Ti-E inverted microscope body with Perfect focus system (Nikon Instruments, Japan). Data was collected in Spinning Disk 40µm pinhole mode on the Zyla 4.2 sCMOS camera using Andor Fusion acquisition software. Image processing and analysis was performed in Fiji/ImageJ. Labelled cilia lengths were measured and plotted in Prism software, where Students t-test statistical analysis was performed.

### Live imaging of retinal slices

Mice were sacrificed by cervical dissection and eyes immediately enucleated. Keratectomy, lensectomy and sclerectomy were performed and retinas embedded in 5% agar and 150 *µ*m sections made using a fully automated VT1200S vibrating blade microtome (Leica Biosystems). Sections were cultured in retinal media (DMEM extra (Sigma) with N2 supplement (Life Tech) and B27 supplement (Life Tech)) in Ibidi 35mm glass bottomed wells at 37 °C in 5% CO2. Slices were incubated with 1 *µ*M SiR-Actin in DMSO (Spirochrome), 10 *µ*M Verapamil in DMSO (Spirochrome) and of 0.6 *µ*M SNAP-Cell TMR-Star in DMSO (New England Biolabs (S9105S) for 120 mins as per manufacturers’ instructions. NucBlue™ Live ReadyProbes™ Reagent (Hoechst 33342; Thermo Fisher) was added before live imaging. Images were acquired using a x60 water immersion lens on the multimodal Imaging Platform Dragonfly (Andor technologies, Belfast UK) equipped with 405, 445, 488, 514, 561, 640 and 680nm lasers built on a Nikon Eclipse Ti-E inverted microscope body with Perfect focus system (Nikon Instruments, Japan). Images were captured in a single z stack every 30 seconds and data collected in Spinning Disk 40µm pinhole mode on a Zyla 4.2 sCMOS camera using Andor Fusion acquisition software. Environmental control of the cultured retinal slices was maintained during imaging with a Okolab bold line stage top incubation chamber incorporating temperature and humidified CO2 control (Okolab S.R.L, Ottaviano, NA, Italy). Live imaging was analysed using Imaris software. Tissue drift was corrected using an autoregressive motion tool by stabilising translational movement of photoreceptor nuclei. Movement of actin microfilaments was tracked by individually selecting the region of interest and measuring Brownian motion over time captured. 6 to 20 mobile actin filaments were analysed per sample. Data was exported to Excel and total distance moved in the X axis over time was calculated. Data for all filaments was then plotted in Prism software, where Student’s t-test statistical analysis was performed.

### Isolation of rod outer segments (ROS)

Mice were maintained in a 12/12 hour light (400 Lux)/dark cycle. Wildtype C57BL/6J, were purchased from Jackson lab (Bar Harbor, ME). 3-month males of RpgrEx3d8 and C57BL/6J mice used for tissue samples were euthanized by CO2 inhalation prior to dissection following American Association for Laboratory Animal Science protocols. Preparation of purified mouse ROS was modified from.(44, 45) Briefly, freshly dissected retinas were collected in 200 *µ*l Ringer’s buffer (10 mM HEPES, 130 mM NaCl, 3.6 mM KCl, 1.2 mM MgCl2, 1.2 mM CaCl2, 0.02 mM EDTA, pH 7.4) with 8% (v/v) OptiPrep (Sigma) under dim red light. Retinas were pipetted up and down with a 200-µl wide orifice tip 50 times, and then centrifuged at 400 × g for 2 min at room temperature. The supernatants containing ROS were collected on ice. The process was repeated 4-5 times. All samples were pooled and loaded onto the top of 10, 15, 20, 25 and 30% (v/v) OptiPrep step-gradient and centrifuged for 60 min at 19,210 × g at 4 °C in a TLS-55 rotor (Beckman Coulter). The ROS band was collected with a 18G needle, diluted with Ringer’s buffer to 3ml, and pelleted in a TLS-55 rotor for 30 min at 59,825 × g at 4 °C. The ROS pellet was resuspended in Ringer’s buffer, and this suspension was applied to grids for cryo-electron microscopy.

### Cryoelectron tomography and image processing

Rod cell fragments containing outer segments, CC, and portions of the IS were collected from WT and *Rpgr*^*Ex3d8*^ mice by iso-osmotic density-gradient centrifugation and applied to EM grids as described above and previously(18, 43, 45). Briefly, isolated WT or *Rpgr*^*Ex3d8*^ cilia were mixed with BSA-stabilized 15 nm fiducial gold (2:1), and 2.5-3 *µ*l of the mixture was deposited on the freshly glow-discharged Quantifoil carbon-coated holey grids (200 mesh, Au R3.5/1) and blotted from the frontside or backside before plunging frozen in liquid ethane using a Vitrobot Mark IV (FEI, Inc.) or automated plunge-freezing device Lecia EMGP (Lecia, Inc.). The frozen-hydrated WT and *Rpgr*^*Ex3d8*^ specimens were stored in liquid nitrogen before imaging. The frozen-hydrated samples were imaged on a JEOL 3200FSC electron microscope operated at 300 kV using a K2 Summit direct electron detector camera or on a JEOL JEM2200FS operated at 200 KV microscope with a DE12 direct electron detector. Both microscopes were equipped with a field emission gun, an in-column energy filter. Single-tilt image series were automatically collected using SerialEM(46) at a defocus range of 8-12 *µ*m and a magnification of 12,000x (equivalent to 3.6 Å/pixel, WT) and 15,000x (equivalent to 4.2 Å)/pixel, KO) on each microscope. The total electron dose per tomogram was 80–100 electrons/Å2, as typical for cellular cryoET. Each tilt series, for WT, has 35 tilt images covering an angular range 51° to +51° with 3° increment (±51°, 3° increment); for *Rpgr*^*Ex3d8*^, there were 51 tilt images with 2° increment (±50°, 2° increment). Tilted images were automatically aligned and reconstructed using the automated workflow in EMAN2 software.(47) Tomographic reconstructions and 3D surface rendering of sub-tomogram averages were generated and visualized using IMOD(48) and UCSF Chimera(49). For the segmentation, to enhance the contrast, tomograms were averaged by two (bin2) or four (bin4) voxels and filtered uniformly using a low-pass filter (set up at 80 Å) to reduce the noise. Structural features such as actin filaments, MT and PM were manually annotated using IMOD(50, 51) and Scripts from EMAN2(52) software package. The segmented maps were visualized using UCSF Chimera 3D software package.

### Subtomogram averaging of actin filaments

The actin filaments were picked from 4x binned tomograms before finer alignment with 1x binned data. Approximately 700 volumes of 60 × 60 × 60 nm actin filaments were extracted at the base of ROS region from WT and *Rpgr*^*Ex3d8*^ tomograms. The initial models (low-pass filtered to 60 Å) were generated using EMAN2 routine(53–55) without applying any symmetry. The subsequent iterative subtomogram refinement using EMAN2(47) yielded the averaged structures of actin filaments at 30 Å resolution. The refinement was performed in ‘gold-standard’ fashion with all particles randomly split into two subsets with resolution measured by the Fourier shell correlation of the density, using the ‘gold’ standard cut-off criterion of FSC=0.143.For the WT samples, approximately 400 volumes of 60 × 60 × 60 nm actin filaments were extracted at the base of ROS region from five tomograms. For *Rpgr*^*Ex3d8*^, approximately 350 volumes of 60 × 60 × 60 nm actin filaments were extracted at the base of ROS region from 6 tomograms. The previously reported F-actin periodicity of to 5.9 nm was observed in initial models for both WT and RpgrEx3d8. The results obtained with these initial models were consistent with the results obtained when using previous F-actin models (with 37 nm, and 27 nm half helical repeat lengths and low-pass filtered to 60 Å)(56) as initial references to initiate iterative refinement of subtomogram-averaged models.

### Quantification and statistical analysis

All statistical analysis was carried out using GraphPad Prism 8 (version 8.4.1; GraphPad software, USA) as described in the text. To determine statistical significance, unpaired t-tests were used to compare between two groups, unless otherwise indicated. The mean ± the standard error of the mean (SEM) is reported in the corresponding figures as indicated. Statistical significance was set at P<0.05.

## Supporting information

Megaw Supplemental Figures

## Acknowledgements

We thank Hemant Khanna for providing his RPGR antibody. We acknowledge the proteomic facility at the Duke Eye Centre for carrying out the 6 week old rod outer segments mass spec experiment. We thank Vadim Arshavsky for sharing his PRCD antibody and protocols, access to core facilties and for specialized mass spec analysis. We thank Craig Nicol for his help in making and formatting the movies for this paper. This work was funded by the Wellcome Trust (RM; 219607/Z/19/Z, FM; 215343/Z/19/Z, MJ; ISSF, AvK; 208402/Z/17/Z), the National Institute for Health (AM; F32 EY031574, TGW and ZZ; R01-EY026545, TGW and FH; R01-EY031949), Fight for Sight (FN; 5179 / 5180), the Medical Research Council (PM and LM; MC-UU-12018/26), Cancer Research UK (LMM; A24452) and the Welch Foundation (TGW; Q-0035).

## Contributions

RM and PM conceived the project. RM, AM, ZZ, LCM, LMM, TGM and PM designed the experiments. RM, AM, ZZ, FN, FM, LCM, AvK, LM, FH and MKJ performed the experiments. RM, AM, ZZ, FN, FM, LCM, AvK, TGM and PM analysed the data. RM, LMM, TGM and PM coordinated the study and provided guidance. RM, AM, LMM, TGM and PM wrote the paper. All of the authors discussed the results and approved the final version of the manuscript.

