## Supplementary material for "Ciliary tip actin dynamics regulate the cadence of photoreceptor disc formation": Megaw Supplemental Figures

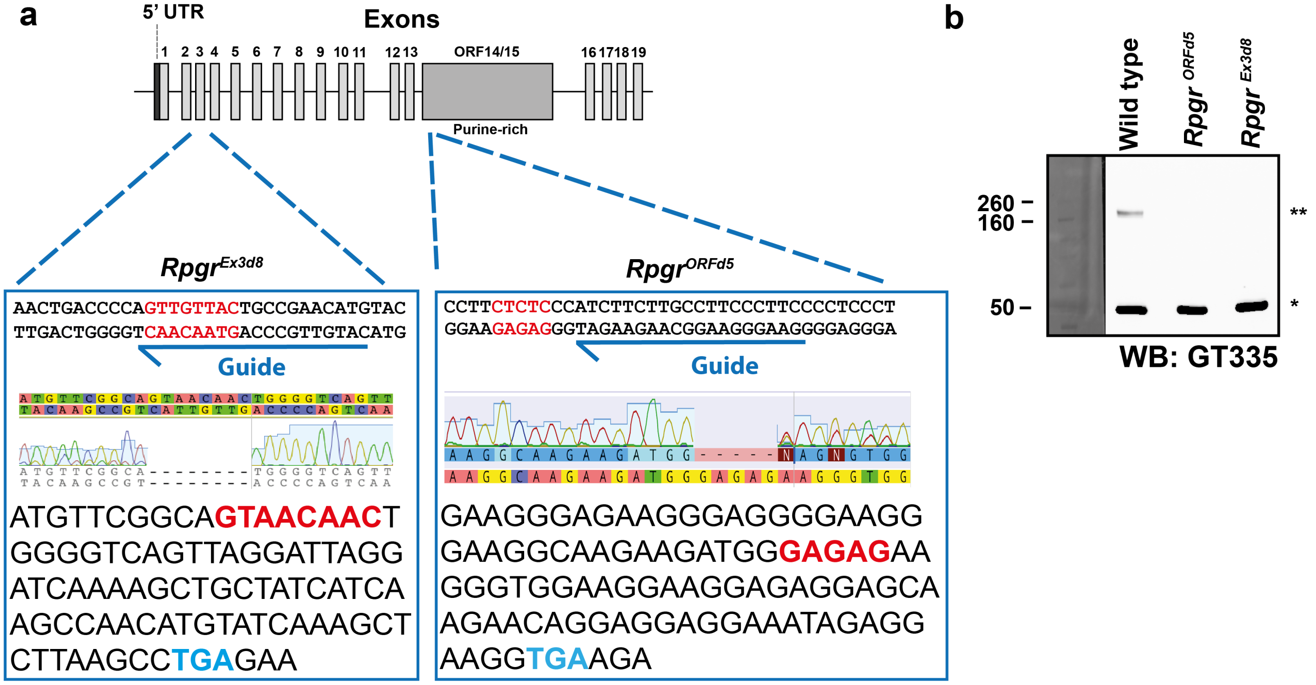


**Extended Data Figure 1. Generation of novel murine *Rpgr* disease models.** **(a)** Targeted pronuclear injection of CRISPR/Cas9 products using guide RNAs as indicated (see methods) induced deletions (red text) in Exon 3 (bottom left panel; termed *Rpgr*$^{Ex3d8}$) and the open reading frame (bottom right panel; termed *Rpgr*$^{ORFd5}$) of *Rpgr*, as detected by Sanger sequencing. This resulted in premature termination codons being shifted into frame (blue text). **(b)** Immunoblotting using a GT335 antibody detects the polyglutamylated, retinal specific isoform of Rpgr (**; see wild type lane)(32). This band was lost in both the *Rpgr*$^{Ex3d8}$ and *Rpgr*$^{ORFd5}$ models. The GT335 antibody also labels polyglutamylated tubulins (*), which serve as a loading control.

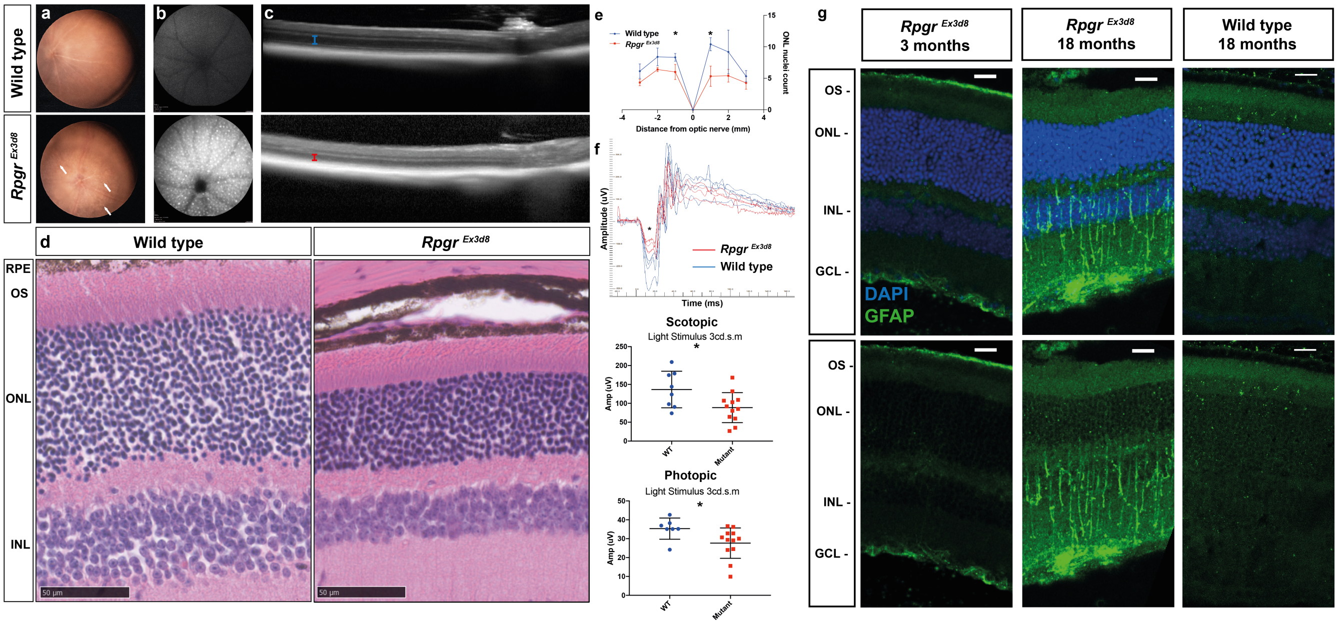


**Extended Data Figure 2. A mouse model of *RPGR/XLRP*, *Rpgr*$^{Ex3d8}$, undergoes loss of photoreceptor function and structure.** **(a)** Fundoscopy shows 18 month *Rpgr*$^{Ex3d8}$ mouse develops white punctate retinal lesions (white arrows). **(b)** Blue light autofluorescence shows accumulation of hyperautofluorescent material in *Rpgr*$^{Ex3d8}$ retina. **(c)** *In vivo* optical coherence tomography shows outer nuclear layer (photoreceptor) thinning in *Rpgr*$^{Ex3d8}$ mice at 18 months (blue bracket in wild type; red bracket in mutant). **(d)** Hematoxylin and eosin staining of 18 month mice supports *in vivo* imaging, with thinning of ONL in *Rpgr*$^{Ex3d8}$ mouse (RPE = retinal pigment epithelium; OS = outer segment; ONL = outer nuclear layer; INL = inner nuclear layer). **(e)** Spider plot demonstrates loss of ONL (n = 3 per experimental group;* = < 0.05). **(f)** Electroretinogram shows loss of retinal function in *Rpgr*$^{Ex3d8}$ mice; top panel shows representative scotopic tracings, with flattening of electronegative downflection ('a' wave; *) in *Rpgr*$^{Ex3d8}$ mice, signalling loss of rod function; middle panel shows a-wave amplitudes in response to 3 candela dark-adapted stimulation; bottom panel shows amplitude of light-adapted flicker response, signalling loss of cone function (n = 7-12 per experimental group; * = < 0.05). **(g)** Retinal stress occurs in *Rpgr*$^{Ex3d8}$ retina, as evidenced by increased glial fibrillary acidic protein (GFAP) immunolabeling throughout radial length of Müller cells at 18 months. Of note, GFAP upregulation is not seen at 3 months, when *Rpgr*$^{Ex3d8}$ mouse has already developed abnormal outer segment architecture (see Figs. 2 a,b). (Scale bars; d = 1 mm; g = 20 $\mu$m)

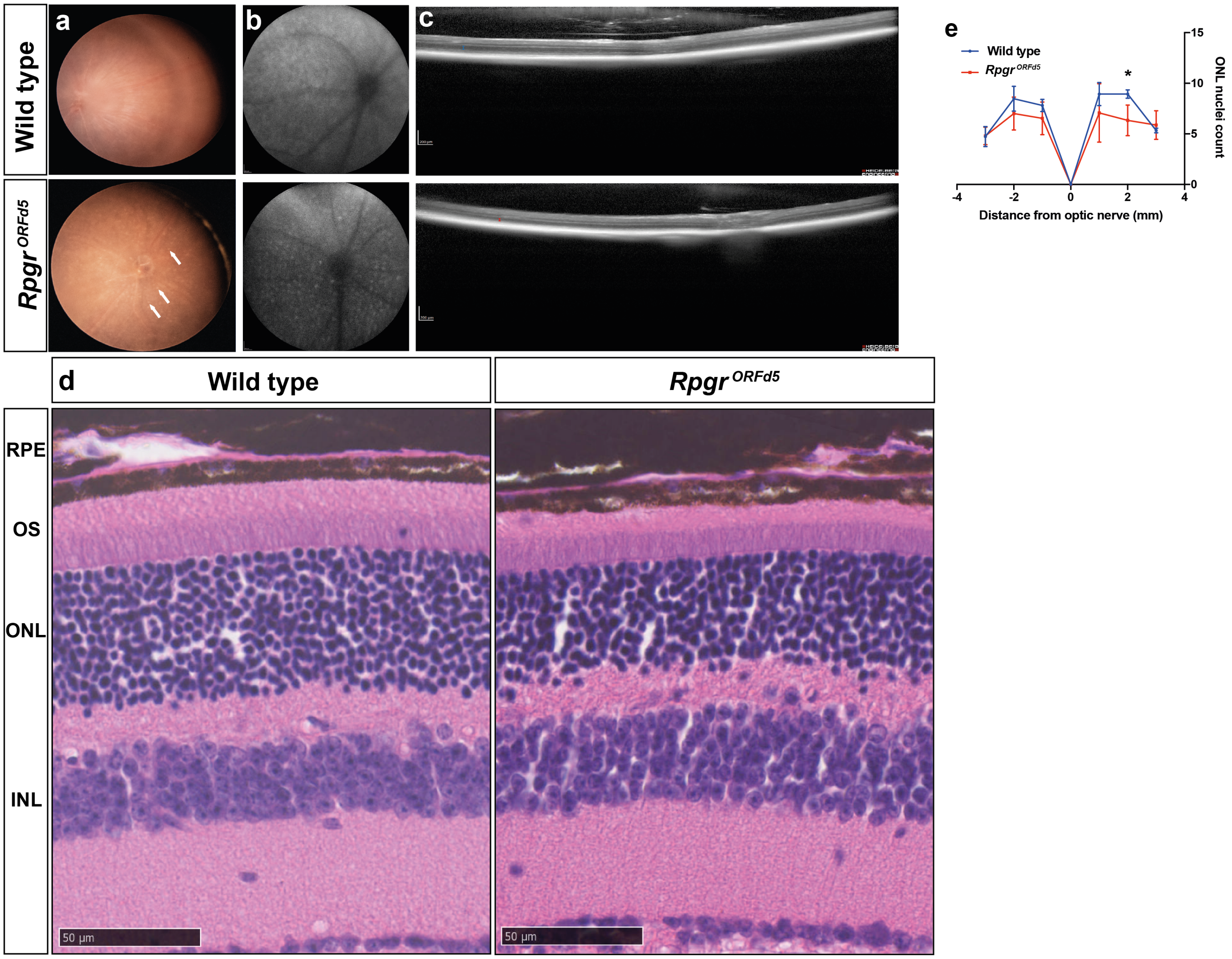


**Extended Data Figure 3. A mouse model of *RPGR/XLRP*, *$Rpgr^{ORFd5}$*, undergoes loss of photoreceptor function and structure.** (**a**) Fundoscopy shows *$Rpgr^{ORFd5}$* mouse develops white punctate retinal lesions (white arrows). (**b**) Blue light autofluorescence shows accumulation of hyperautofluorescent material in *$Rpgr^{ORFd5}$* retina. (**c**) In vivo optical coherence tomography shows outer nuclear layer (photoreceptor) thinning in *$Rpgr^{ORFd5}$* mice at 18 months of age (blue brackets in wild type; red brackets in mutant). (**d**) Conventional histology of 18 month old mice supports in vivo imaging, with thinning of outer nuclear layer in *$Rpgr^{ORFd5}$* mouse (RPE = retinal pigment epithelium; OS = outer segment; ONL = outer nuclear layer; INL = inner nuclear layer;). (**e**) Spider plot demonstrates loss of outer nuclear layer across the retina (n = 3 per experimental group; * = < 0.05). (Scale bars; c = 200 $\mu$m; d = 1 mm).

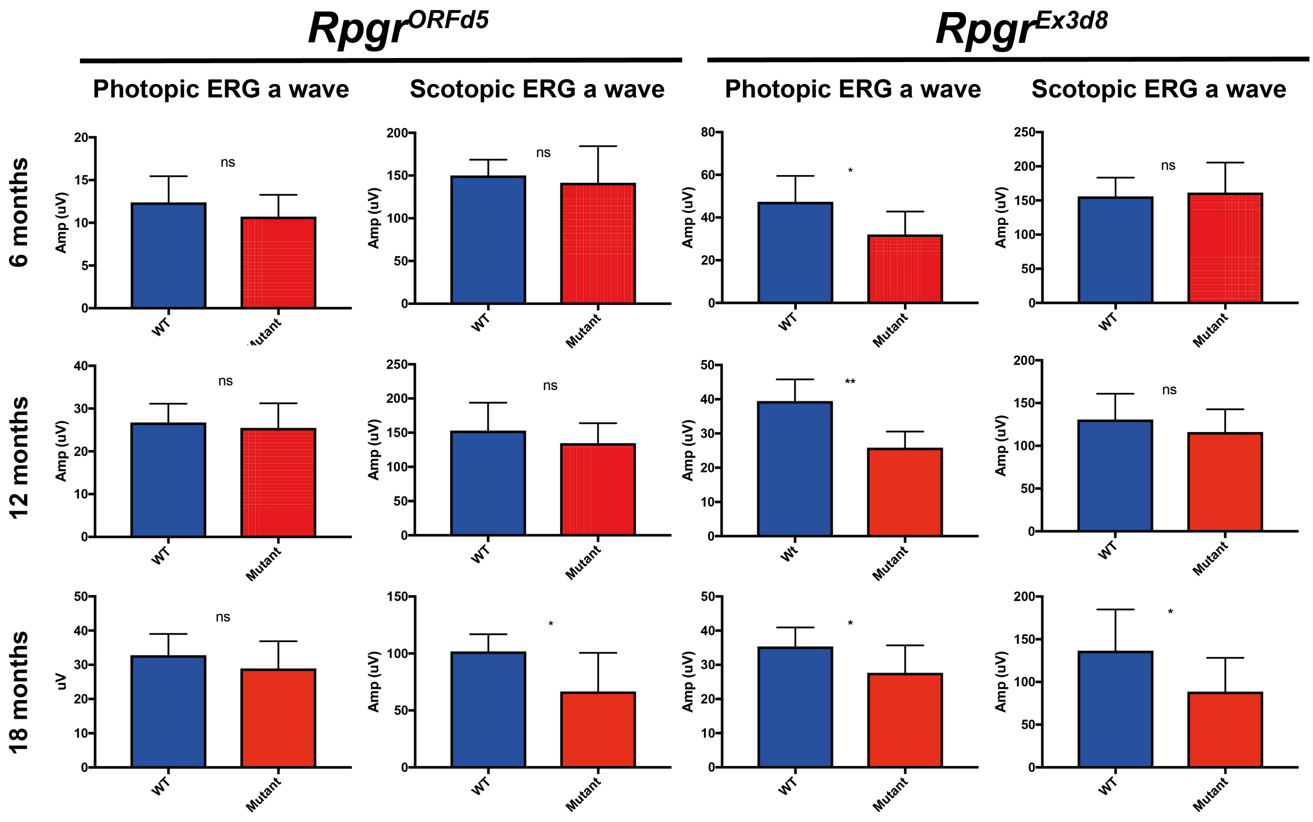


**Extended Data Figure 4. Temporal ERG data shows retinal function declines over time in both mouse models of *Rpgr* disease.** Analysis of *Rpgr*$^{ORFd5}$ (left 2 columns) and *Rpgr*$^{Ex3d8}$ (right 2 columns) mice at 6, 12 and 18 months shows loss of rod photoreceptor (scotopic light stimulus 3cd.s.m $^{-2}$) function at 18 months in both strains. Analysis of cone photoreceptor (photopic light stimulus 3cd.s.m $^{-2}$) function shows early loss of cone function in *Rpgr*$^{Ex3d8}$ mice but not in *Rpgr*$^{ORFd5}$ mice (n = 7-12 per experimental group).

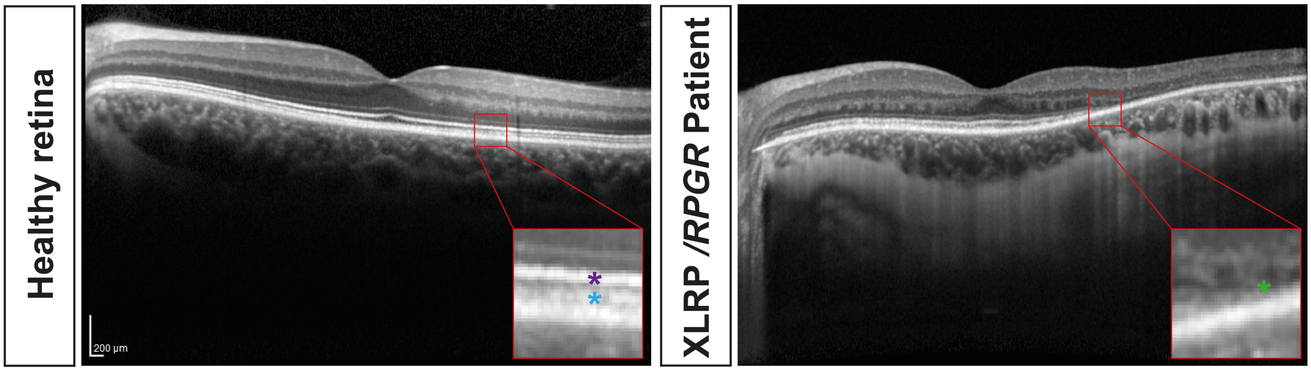


**Extended Data Figure 5. Optical coherence tomography (OCT) shows loss of ellipsoid zone in *RPGR/XLRP* patient.** OCT of healthy volunteer (left panel). Magnified insert shows the ellipsoid zone (magenta star) and interdigitating zone (cyan star) that together represent the photoreceptor outer segment. OCT of patient with RPGR/XLRP (right panel). Magnified insert shows loss of ellipsoid zone temporal to macula (extending centrally from right of scan up to green star).

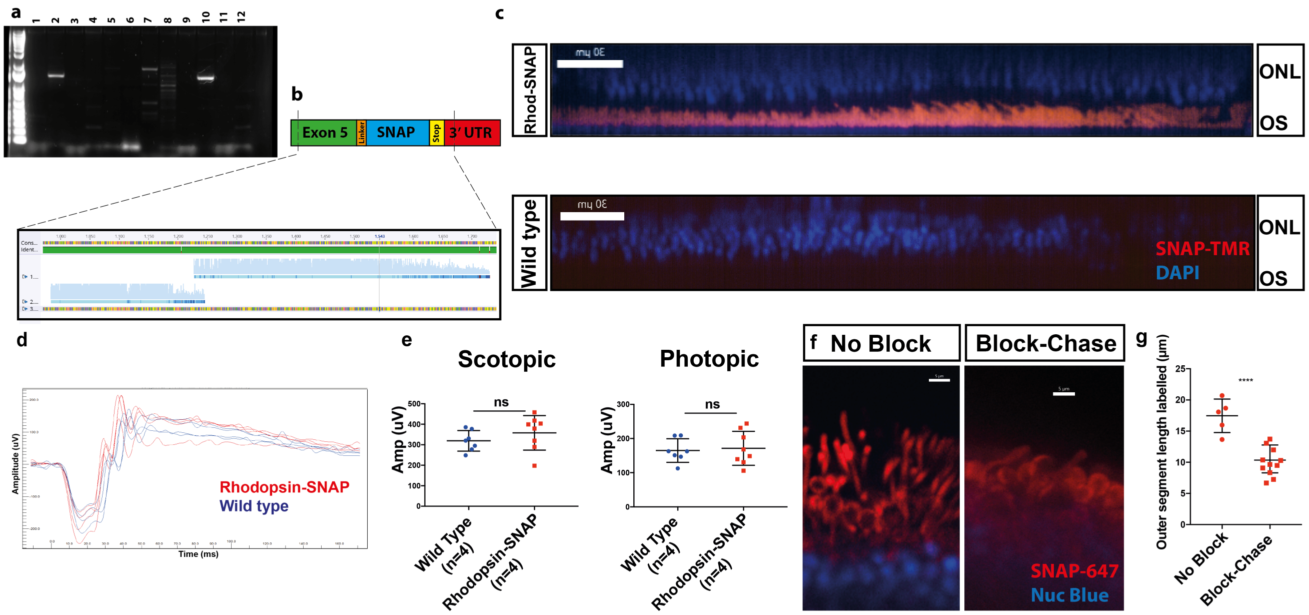


**Extended Data Figure 6. Generation of an outer segment turnover biosensor with an endogenously tagged *Rhod*$^{SNAP}$ mouse.** **(a)** Genotyping of founder mice following pronuclear injection of CRISPR/Cas9 products showed mice 2 and 10 as potential SNAP knock ins. **(b)** Sanger sequencing showed successful in frame knock in of SNAP at the C terminus. **(c)** Intravitreal injection SNAP-TMR shows photoreceptor outer segment labelling in *Rhod*$^{SNAP}$ mouse but not wild type (ONL = Outer Nuclear Layer; OS = Outer Segment. **(d,e)** Electrodiagnostic testing shows no loss of photoreceptor function in *Rhod*$^{SNAP}$ mice at 6 months of age (n = 7-8 per genotype); either for rod function (left graph; scotopic light stimulus 3 cd.s.m$^{-2}$) or cone function (right graph; photopic light stimulus 3 cd.s.m$^{-2}$). **(f)** Incubation of naive (no SNAP block) *Rhod*$^{SNAP}$ retinal slice cultures with fluorescent SNAP ligand shows labelling of entire outer segment (left panel). Incubation of Rho-SNAP retina slice culture with fluorescent SNAP ligand 72 hours after intravitreal treatment with SNAP-block shows shorter labelling of outer segment; consistent with newly formed discs. **(g)** Graph showing length of fluorescently labelled outer segment following retinal slice incubation with and without SNAP-block (n=3 per experimental group; **** = $p<0.0001$) (Scale bars; c = 30 $\mu$m; f = 5 $\mu$m)

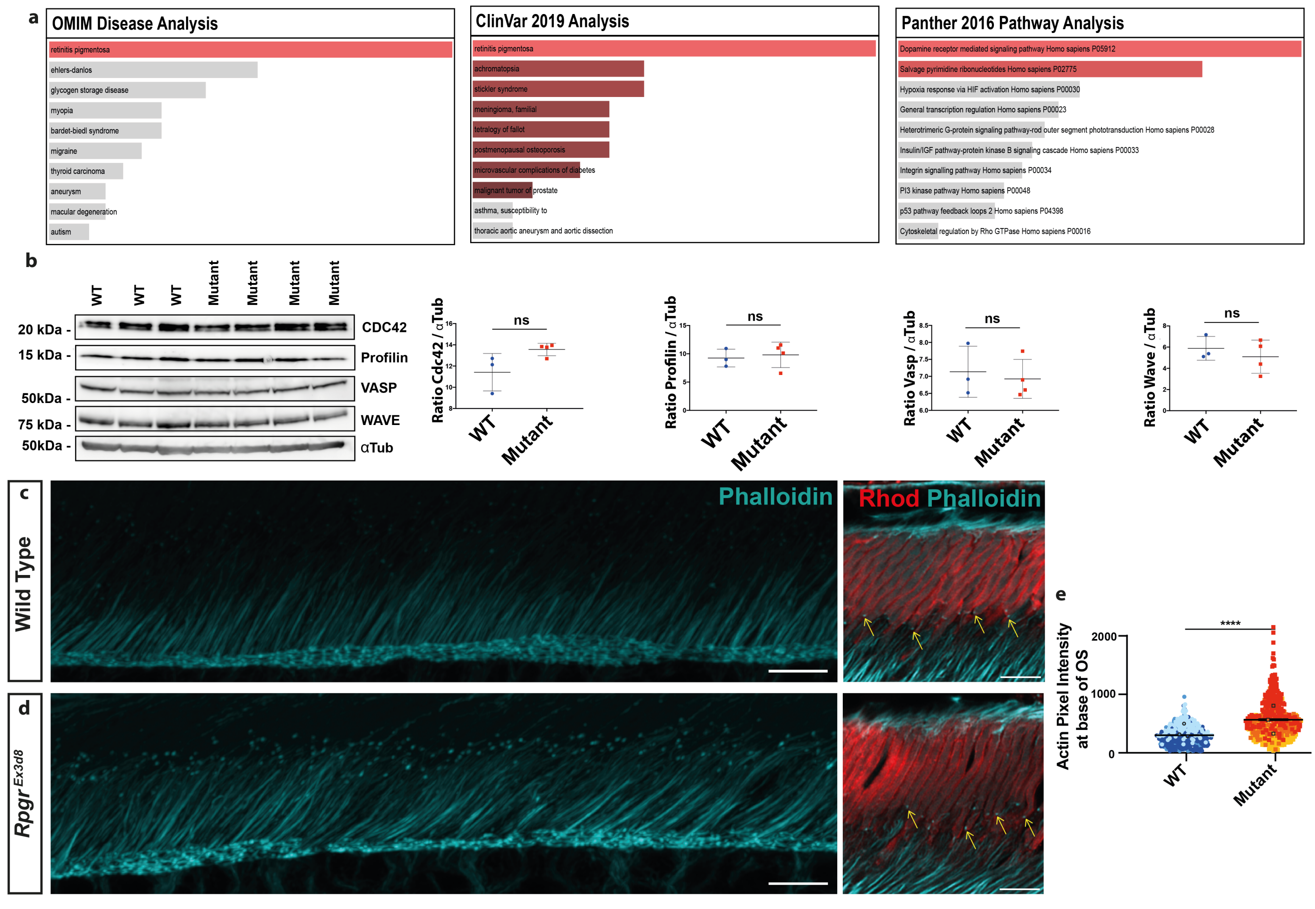


**Extended Data Figure 7. The *Rpgr*$^{Ex3d8}$ mouse has actin dysregulation in the connecting cilium (CC). (a)** Computational analysis of differentially expressed proteins using ClinVar 2019 and OMIM disease libraries ranks retinitis pigmentosa as the top disease in *Rpgr*$^{Ex3d8}$ retina compared to wild type control. Comparison of differentially expressed proteins to Panther 2016 protein pathway database ranked cytoskeletal regulation by Rho GTPases as a significantly disrupted pathway. **(b)** Immunoblotting of various actin binding proteins and nucleators show no significant difference in protein levels in *Rpgr*$^{Ex3d8}$ retina lysates compared to wildtype (y axis denotes ratio of protein of interest to $\alpha$-Tubulin loading control). Airyscan comparison of wild type ((**c**); top panels) and *Rpgr*$^{Ex3d8}$ ((**d**); bottom panels) retina shows increased actin polymerisation in the distal photoreceptor CC in *Rpgr*$^{Ex3d8}$ mice, as labelled by phalloidin. Right panels, at higher magnification, depict localisation of actin puncta (yellow arrows) within the CC, at the base of the rhodopsin-positive outer segment. **(e)** Graph showing sum pixel intensity of photoreceptor CC phalloidin staining (n=3 animals per genotype; **** = $p<0.0001$) (Scale bars = 5 $\mu$m).

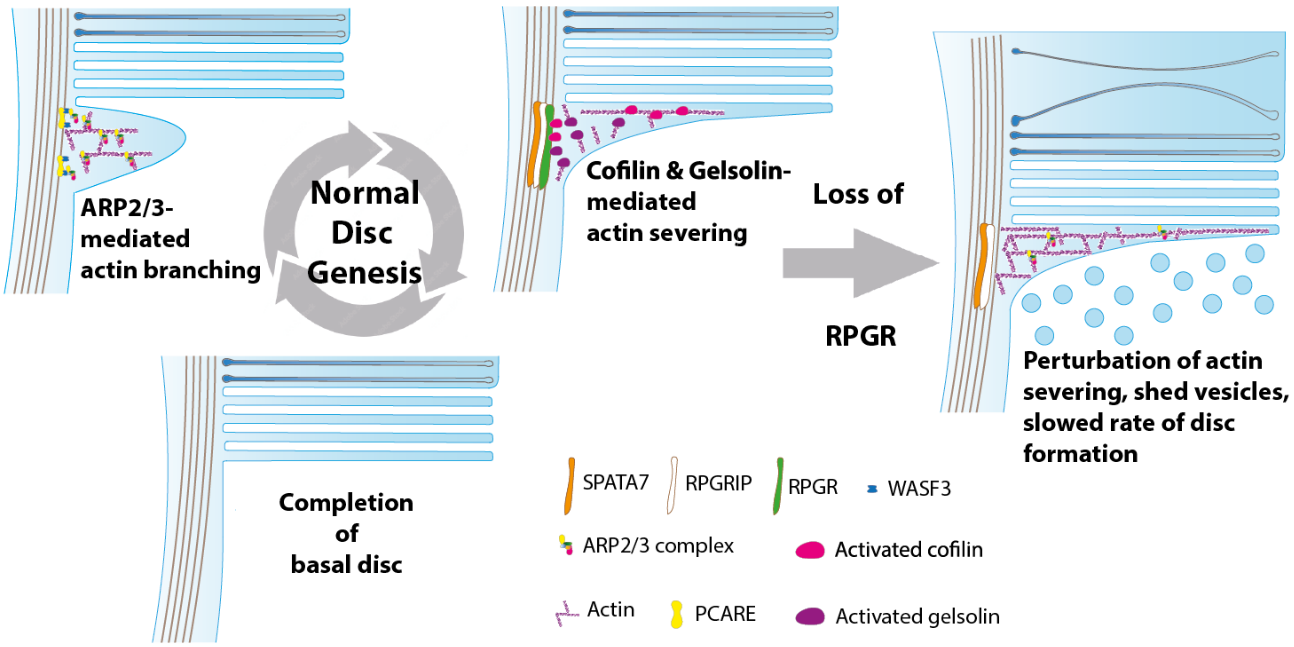


**Extended Data Figure 8. RPGR is essential for controlling the cadence of outer segment disc genesis.** Photoreceptor disc formation is an active, actin driven process. PCARE and WASF3 mediate ARP2/3-driven actin filament polymerisation, resulting in cilia membrane deformation and disc initiation. Membrane addition leads to disc elongation until SPATA7, RPGRIP and RPGR regulate gelsolin- and cofilin-mediated actin severing, allowing disc formation to complete. Loss of RPGR perturbs this actin severing, with abnormal discs instead being aborted and shed as ectosome-like vesicles. The resulting slowed rate of disc formation causes retinal stress, photoreceptor degeneration and loss of vision.

**Extended Data Movie 1. Live imaging of retinal slice cultures captures actin dynamics in the photoreceptor connecting cilium.** Spinning disc microscopy of live retinal slices depicting photoreceptor nuclei (NucBlue probe; blue), actin (SiR-actin; magenta) and outer segments (SNAP-TMR; cyan) over time.

**Extended Data Movie 2. High magnification view comparing actin dynamics in wild type and emphRpgr$^{Ex3d8}$ connecting cilia.** Imaris software tracks mobile actin filaments (projected as coloured 'dragon tails' in movie) in Wild Type (left panel) and emphRpgr$^{Ex3d8}$(right panel) connecting cilia. Total distance moved of mobile filaments in the x plane over time imaged was then calculated (see **Figure 6**).
